# Benchmarking brain organoid recapitulation of fetal corticogenesis

**DOI:** 10.1101/2022.04.22.488753

**Authors:** Cristina Cheroni, Sebastiano Trattaro, Nicolò Caporale, Alejandro López-Tobón, Erika Tenderini, Flavia Troglio, Michele Gabriele, Raul Bardini Bressan, Steven M Pollard, William T Gibson, Giuseppe Testa

**Affiliations:** Department of Experimental Oncology, European Institute of Oncology IRCCS, Via Adamello 16, 20139, Milan, Italy; Department of Oncology and Hemato-Oncology, University of Milan, Via Santa Sofia 9, 20122, Milan, Italy; Human Technopole, Viale Rita Levi-Montalcini 1, 20157, Milan, Italy; Telethon Institute of Genetics and Medicine (TIGEM), Pozzuoli, Naples, Italy, Via Campi Flegrei, 34 - 80078; Department of Biological Engineering, MIT, Cambridge, MA; Broad Institute of MIT and Harvard, Cambridge, MA; Centre for Regenerative Medicine, Institute for Regeneration and Repair, and Edinburgh Cancer Research UK Centre, University of Edinburgh, Edinburgh, UK. EH16 4UU; BC Children’s Hospital Research Institute, 950 West 28th Avenue, Vancouver, BC, Canada V5Z 4H4; Department of Medical Genetics, University of British Columbia, Vancouver, BC, Canada V6T 1Z4

**Keywords:** Brain organoids, human corticogenesis, neurodevelopment, disease-modelling, transcriptomics, neurodevelopmental disorders, psychiatric disorders, brain development, gene co-expression patterns

## Abstract

Brain organoids are becoming increasingly relevant to dissect the molecular mechanisms underlying psychiatric and neurological conditions. The *in vitro* recapitulation of key features of human brain development affords the unique opportunity of investigating the developmental antecedents of neuropsychiatric conditions in the context of the actual patients’ genetic backgrounds. Specifically, multiple strategies of brain organoid (BO) differentiation have enabled the investigation of human cerebral corticogenesis *in vitro* with increasing accuracy. However, the field lacks a systematic investigation of how closely the gene co-expression patterns seen in cultured BO from different protocols match those observed in fetal cortex, a paramount information for ensuring the sensitivity and accuracy of modeling disease trajectories. Here we benchmark BO against fetal corticogenesis by integrating transcriptomes from in-house differentiated cortical BO (CBO), other BO systems, human fetal brain samples processed in-house, and prenatal cortices from the BrainSpan Atlas. We identified co-expression patterns and prioritized hubs of human corticogenesis and CBO differentiation, highlighting both well-preserved and discordant trends across BO protocols, including different degrees of heterochronicity in differentiation across BO models compared to fetal cortex. Our approach provides a framework to directly compare the extent of *in vivo/in vitro* alignment of neurodevelopmentally relevant processes and their attending temporalities, structured as an accessible resource to query for modelling human corticogenesis and the neuropsychiatric outcomes of its alterations.

## INTRODUCTION

The introduction of cell reprogramming technologies and 3D brain organoids (BO) have made the spatial and temporal dynamics of human brain development experimentally accessible. BO are thus becoming central to the investigation of how genetic vulnerabilities or environmental perturbations can alter physiological neurodevelopment and seed the unfolding of psychiatric and neurological conditions^1,2^. As we and others recently demonstrated, the exposure of specific windows of vulnerability is proving of particular value, highlighting how temporally defined or even transient alterations in neurodevelopmental trajectories can bring about major mental health outcomes, from language delay to ASD^3–5^. Indeed, a growing body of literature testifies to the edge that BO are bringing to the modelling of complex neuropsychiatric disorders, from autism spectrum disorder to schizophrenia, bipolar disorder and beyond^6–8^. Given the central role of the cortex for the higher order functions primarily affected in neuropsychiatric conditions, determining how closely BO recapitulate human cerebral corticogenesis is thus crucial for harnessing their full potential as *in vitro* models of neurodevelopmental processes and their physiopathological outcomes. To this end, comparison between ex-vivo human fetal cortex and BO gene expression has started to uncover the extent of this recapitulation, alongside the peculiarities of different methods. Specifically, the transcriptomic and epigenomic landscapes of BO were characterized and compared to isogenic fetal cortices, finding significant overlaps^9^. Very long-term BO cultures were meanwhile shown to capture early post-natal developmental transitions^10^, while spatial similarity maps of BO against reference were generated to assess organoid engineering protocols and to annotate cell fates^11^. Further detail emerged from integrative analyses showing an overexpression of extracellular matrix (ECM)-related genes in BO associated with the first steps of differentiation in 2D^12^, a higher vulnerability to cellular stress in some organoid cultures compared to primary tissue^13,14^, and the existence of protocol-specific transcriptional bypasses in BO differentiation^15^.

These efforts have provided a wealth of information, which remains however difficult to harmonize and use productively due to the lack of dedicated resources where transcriptional hubs of development/differentiation are categorized, ranked, and made available for consultation. Such tools are needed to help researchers select protocols and time-points based not only on the expression of genes of interest but also on the broader context of functional partitions and temporal dynamics of that expression. Building on these considerations, we benchmarked selected BO paradigms against human corticogenesis with the aim of templating such a resource through a framework that allows its iterative adaptation and growth as the neural modelling field continues to mature. Specifically, we profiled in-house a cohort of cortical BO (CBO)^16,17^, derived from multiple individuals and differentiation rounds over 200 days, and integrated it with i) our in-house cohort of primary samples, ii) publicly available transcriptomic data from BO different for degree of patterning, and iii) prenatal cortical samples of the BrainSpan Atlas (BS). The characterization of the gene expression landscape of BS and CBO led to the definition of co-expression patterns relevant for human prenatal corticogenesis and CBO differentiation, as well as to the ranking and categorization in functional domains of their transcriptional hubs. Development/differentiation co-expression patterns were made available. We then cross-compared CBO differentiation, primary samples, and other BO systems^12,18,19^. These analyses revealed the overlap between co-expression patterns of prenatal cortex and CBO, allowed their visualization in the selected BO systems, and pointed at partial heterochronicity in the transcriptional recapitulation of corticogenesis by different BO methods.

In sum, our analyses represent a benchmark for the interrogation of the transcriptional networks characterizing human prenatal corticogenesis and CBO differentiation and uncover their modulation also in other BO protocols of widespread use for neuropsychiatric disease modelling. By comparing co-expression patterns of human corticogenesis and BO differentiation across protocols, our work represents a template for the benchmarking of BO and their applications in disease modeling.

## RESULTS

### Gene co-expression analysis highlights the transcriptional programs of the prenatal human corticogenesis

To categorize the transcriptional hubs of the developing human cortex, we took advantage of the BrainSpan Atlas BS, the most comprehensive transcriptional characterization of the fetal human brain.

In a landmark effort, Miller et al. profiled by microarray fetal brain at mid-gestation^20^, establishing a first reference transcriptional atlas. More recently, a multi-modal characterization of human brain was performed^21^. We applied a similar approach at the transcriptome level, focusing our effort specifically on the prenatal cerebral cortex. This allowed us to reconstruct the transcriptional circuitries of the human fetal corticogenesis with increased resolution. We selected a total of 162 data points (**Figure 1A**) from cerebral cortex at post-conceptional weeks (PCW) 8-37.

**Figure 1:**
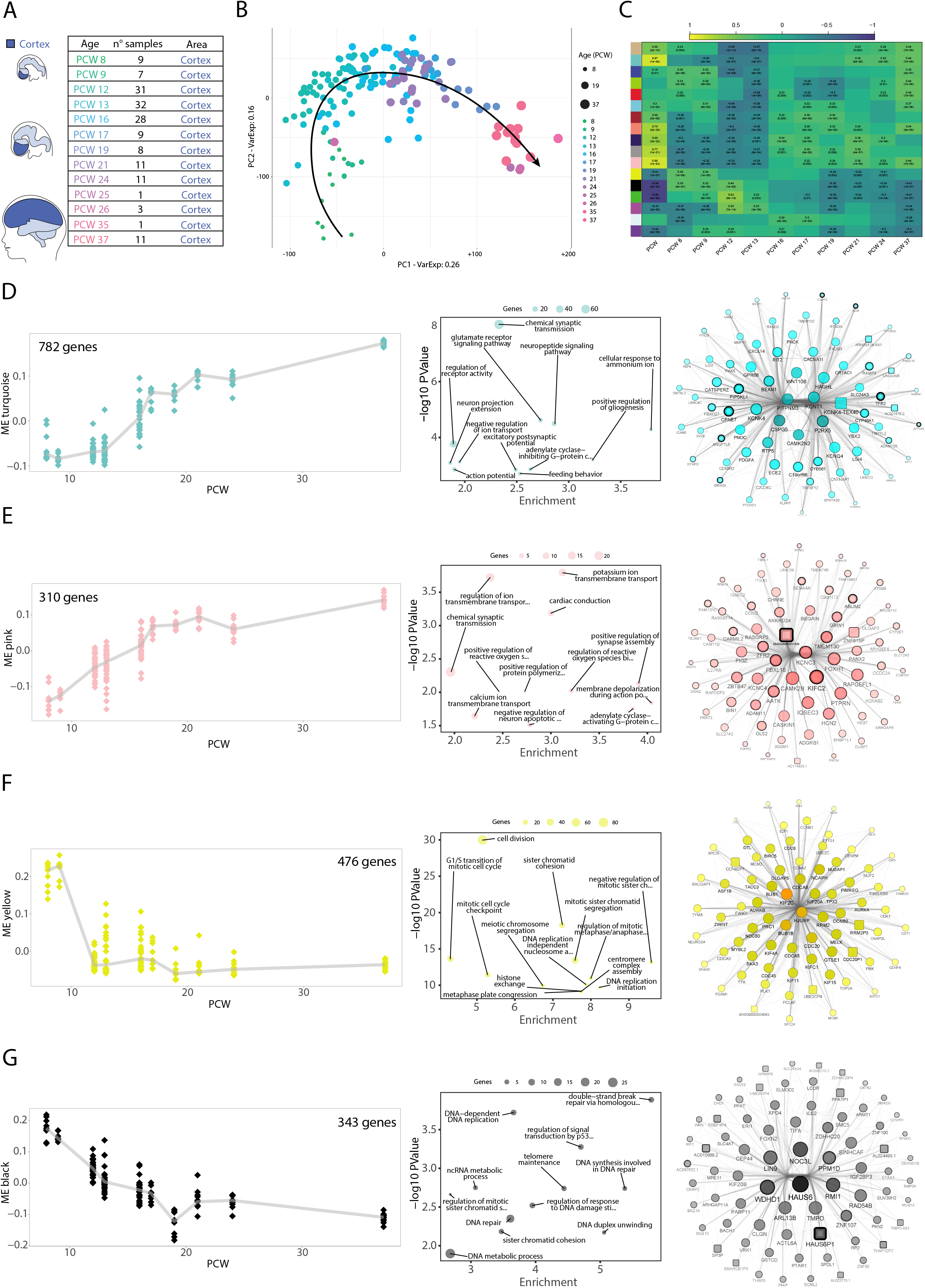
Reconstruction of transcriptional programs of the developing fetal cortex. **(A)** Cohort of fetal cortical samples from the BS Atlas. The number of specimens for each post-conceptional week (PCW) is reported, for a total of 162 bulk RNASeq data points. **(B)** PCA on prenatal cortical samples from BS. Dot color and size is set according to developmental stage. **(C)** WGCNA pinpoints gene modules in the developing cortex. The heatmap shows the correlation between the first principal component of each module (module eigengene, ME) and the developmental stage, either as a continuous variable (PCW) or a categorical variable for each stage. Coefficients of correlation were calculated using Spearman correlation; p-values are reported for significant correlations (p-value < 0.01). Each row represents a gene module and it is identified as with a specific color. (**D-G**) Characterization of BS_Turquoise, BS_Pink, BS_Yellow and BS_Black modules. For each module, ribbon plots visualize the behaviour of the ME (Y-axis) through developmental stages; each dot represents a data point, while the line connects the median value for each PCW. The bubble plots show the p-value (Y-axis) and enrichment score (X-axis) for the top-12 GO categories (ranked according to p-value, Biological Process domain of the ontology); dot size represents the number of module genes belonging to the GO term. Network reconstruction for the top-75 genes of each module, selected according to the intramodular connectivity. Degree, closeness, betweenness and eigenvector are represented by node label transparency, node colour darkness, node border width, and node size/node label font size, respectively. Node shape represents gene biotype, with protein coding genes as circles and non-coding genes as squares.

Principal component analysis (PCA) identified developmental stage as the main driver of sample differences, with PC2 separating very early stages (PCW 8-9) from early ones (PCW 12-13) followed by a progression of time-points in PC1 (**Figure 1B**). Likewise, stage-wise correlation analysis detected a first shift in the transcriptional landscape from PCW 8-9 to 12-24 and a second, less clear-cut, characterizing PCW 25-37 (**Supplementary Figure 1A**). The same analysis on postnatal cortex showed higher homogeneity compared to prenatal time-points (**Supplementary Figure 1B)**, indicating a more profound transcriptional evolution during the fetal phase, especially till PCW24.

We applied gene ontology enrichment analysis (GO) to the top 600 genes associated with PC1 or PC2 according to PCA loadings. We retrieved for PC1 categories associated with ion channels, lipid metabolism and transcriptional regulation, while we found terms related to neuronal maturation and cell division for PC2 (**Figure 1B; Supplementary Figure 1C-D**).

To identify the transcriptional programs regulating corticogenesis in an unsupervised manner, we applied weighted gene co-expression network analysis (WGCNA), uncovering 17 gene co-expression modules (**Figure 1C and Supplementary File 1**). We summarized the co-expression profile of each module by its first PC (module eigengene, ME) and related it to developmental stage, highlighting several modules as positively or negatively correlated. BS_Turquoise, BS_Pink, BS_Grey60 and BS_Midnightblue showed positive correlation, while BS_Black and BS_Green were negatively associated with developmental progression (**Figure 1C**). Other modules displayed changes in more restricted time windows (e.g. BS_Yellow, BS_Blue, BS_Red, **Figure 1C**). The functional characterization of BS modules pointed to clear-cut biological domains for several of them, such as glutamatergic transmission and synapse for BS_Turquoise, ion channels for BS_Pink, DNA replication for BS_Black, and cell division for BS_Yellow (**Figure 1 D-G**). Finally, we reconstructed the co-expression network selecting the top-75 genes (according to intramodular connectivity) and then applied network analysis. Central nodes of BS_Turquoise and BS_Pink networks were related to neuronal functions and included synaptic proteins (CAMK2N2, CAMK2B and GDA), receptor subunits (GRIN1, GRIN3A) and potassium channel subunits (KCNT1, KCNQ4, KCNC3, KCNC4). BS_Midnight-blue included the upper layer neuron markers CUX2 and SATB2 among its most central genes (**Supplementary figure 1F**). BS_Yellow and BS_Black hub genes were strongly enriched in cell cycle genes.

The reconstructed networks also encompassed genes less studied in corticogenesis (e.g. for BS_Turquoise CATSPERZ, HAGHL, ADAMTS8, and KCN4-TEX40), thus allowing us to hypothesize their involvement by a guilt-by-association approach (**Figure 1 D-G and Supplementary figure 1F**).

Overall, we identified transcriptional circuitries related to developmental transitions and functional domains of human corticogenesis, reconstructing relevant gene co-expression networks. These networks, their behaviour, and the relevance of every gene composing them are available in **Supplementary File 1**.

### Cortical brain organoids globally resemble the developing human fetal cortex and evolve in two steps

Upon characterization of the transcriptional dynamics defining human corticogenesis, we investigated the extent of their recapitulation in BO, with an experimental design that allowed us to measure interindividual and technical variability. We analysed in-house differentiated CBO generated as previously described^16^, for a total of 43 single-organoid samples from 4 control hiPSC lines profiled over 200 days. We profiled single organoids to tackle technical variability of differentiation and we analyzed 2 independent organoid batches to measure reproducibility. Additionally, we profiled a cohort of primary fetal CNS tissues (weeks of gestational age -WGA-13 and 15) and 2D cultures (WGA 11, donor 1, and 19, donor 2) for a total of 4 individuals (**Figure 2A**); this dataset, processed as CBO, allowed direct comparisons between the two.

**Figure 2:**
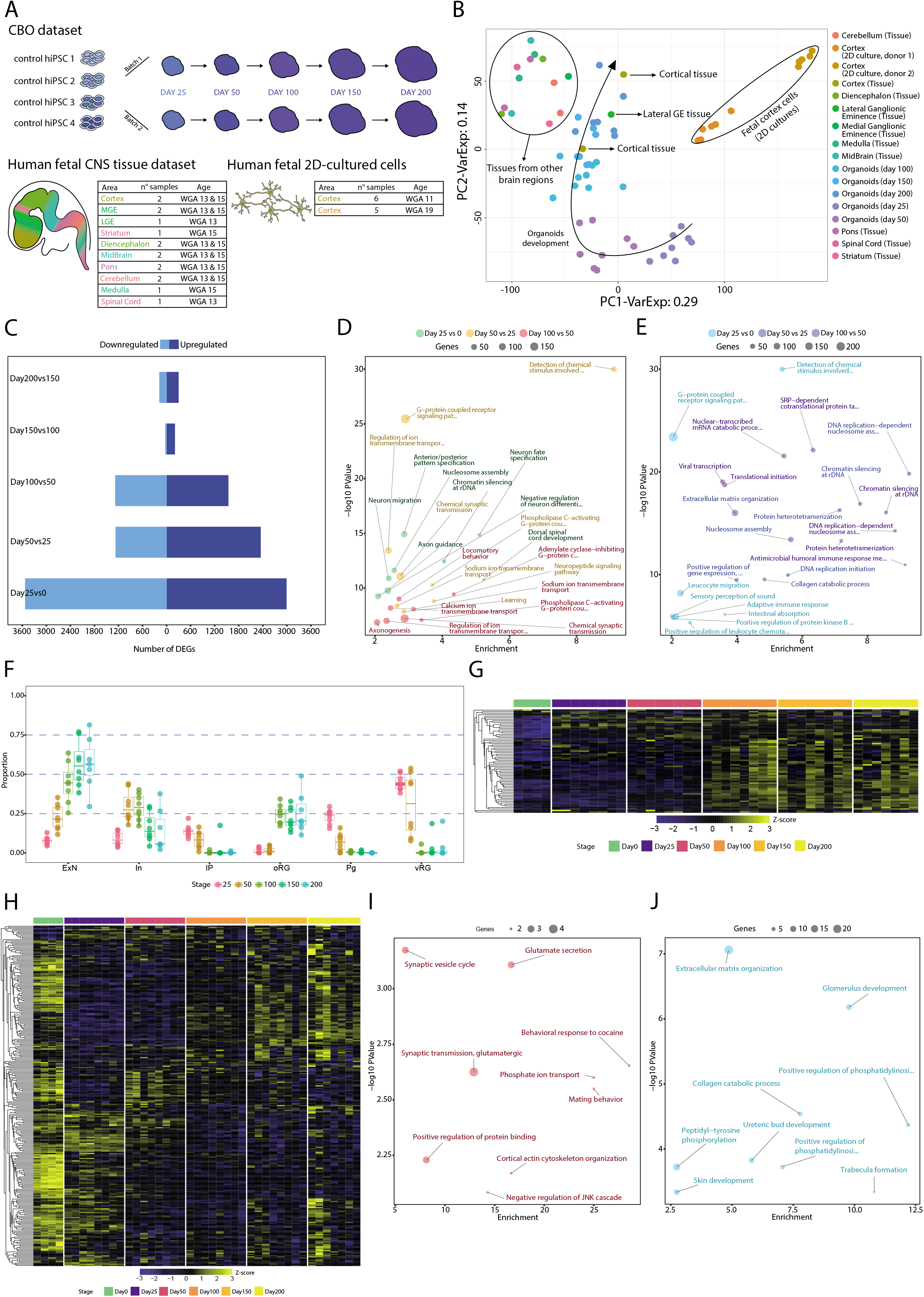
Principal Component and Differential Expression analyses uncovered CBO temporal dynamics. **(A)** Experimental design, detailing the number of individuals, batches and time-points for CBO and number of samples, brain area and WGA for fetal samples. **(B)** PCA on CBO, fetal brain tissues and 2D cultured fetal cortical progenitors, distinguished by dot color. The arrow highlights the distribution of CBO samples throughout stage progression. **(C)** Number of DEGs (FDR < 0.05, log2FC > 1 as absolute value) from stagewise differential expression analysis in CBO dataset. Barplots show the number of up-regulated and down-regulated genes for each comparison. **(D-E)** Bubble plots showing the functional characterization of up-regulated (D) and down-regulated (E) genes by functional enrichment analysis for the following comparisons: (I) Day25 vs Day0; (II) Day50 vs Day25; (III) Day100 vs Day50. The top-8 GO categories from the Biological Process domain of the GO are reported for each comparison. **(F)** Box plots displaying the estimated proportion of excitatory neurons (ExN), inhibitory neurons (In), intermediate progenitors (IP), outer radial glia (oRG), cycling progenitors (Pg), and ventricular radial glia (vRG) from deconvolution in each stage of CBO differentiation. **(G-H)** Heatmaps showing the behaviour of cortex-specific up-regulated (G) and down-regulated (H) DEGs in the CBO dataset. Values are shown as Z-scores calculated on the expression values. **(I-J)** Bubble plots showing the functional characterization of the genes in G and H, respectively.

PCA showed temporal evolution of CBO with early stages forming individual clusters and later time-points resulting more intermingled. CBO evolved towards the fetal tissue, with clustering of mature organoids in proximity of fetal cortex. Conversely, 2D cultures clustered apart (**Figure 2B**). Specific focus on CBO and relative functional analysis allowed us to associate PC1-driving genes to cell cycle and neuronal differentiation (**Supplementary figure 2A-B**).

To quantitatively characterize CBO evolution, we performed stage-wise differential expression analysis (DEA) comparing each stage against the previous. This approach highlighted a biphasic differentiation dynamic, with fast changes until day 100 followed by subtler modulations (**Figure 2C**). The fast-evolving phase was characterized by a large number of differentially expressed genes (DEGs), related to neuronal fate commitment and maturation (upregulated) and cell cycle, transcription, and translation dynamics (downregulated) **(Figure 2D-E)**. The number of DEGs decreased considerably in the slow-evolving phase and confirmed cell division as a prominent down-regulated category. Upregulated genes were more heterogeneous in function **(Supplementary figure 2D-E)**^22–28^.

We then analysed dynamic changes among DEGs by visualizing the fold-change of consecutive comparisons (e.g. day 50vs25 against 100vs50, etc). Neuronal differentiation regulators, such as EOMES, LHX2, and FEZF2^29–31^, were modulated dynamically in the fast-evolving phase, while we detected upregulation of the astrocytic markers HEPACAM, AQP4, AGT and APOE in the slow-evolving phase^32–35^. We also observed upregulation of GABAergic interneuron markers after day 100, in line with other studies^36,37^ (**Supplementary figure 2F**).

We then estimated CBO cell-type proportions with bulk deconvolution exploiting a scRNAseq atlas of the developing human cortex as reference^38^. Deconvolution methods have already been employed to estimate cell type proportions in the fetal brain^21^ and have been shown to give reliable estimates when benchmarked against immunohistochemical quantification of the main adult brain cell types, overcoming some of the selection biases affecting single-cell estimations^39^. From day 25 till day 100 CBO showed increase of excitatory neurons mirrored by a drop in ventricular radial glia, cycling progenitors, and intermediate progenitors. Outer radial glia increased from day 100 onwards (**Figure 2F**).

Lastly, we identified cortex-enriched genes by comparing the cortical samples of our in-house fetal brain tissue dataset (WGA 13,15) against hiPSCs and subsequently excluding common DEGs found with the same approach for other brain areas. We analyzed expression of cortex-enriched DEGs in CBO and found upregulation of genes related to glutamatergic neuron function alongside downregulation of ECM genes (**Figure 2G-J**).

### WGCNA identifies cortical brain organoids’ differentiation trajectories towards the glutamatergic fate

To identify transcriptional patterns in CBO differentiation we applied WGCNA, identifying 14 co-expression modules (**Supplementary Figure 3A, Supplementary File 2**). Correlation with differentiation stage uncovered as strongly correlated modules the CBO_Turquoise and CBO_Black (positive correlation) and CBO_Brown and CBO_Blue (negative correlation) (**Figure 3A**). We analyzed the behaviour of CBO_Turquoise and CBO_Black ME in the 8 samples across time-points (**Figure 3B**) detecting a steady increase over time. While CBO_Turquoise modulation was reproducible across replicates, the CBO_Black was more variable, pointing to a small set of genes (180 genes against the 3279 of the CBO_Turquoise) with a less robust behaviour. The variability of this subset of genes was observed mainly in 2 samples, which however showed patterns in line with the other samples for other modules.

**Figure 3:**
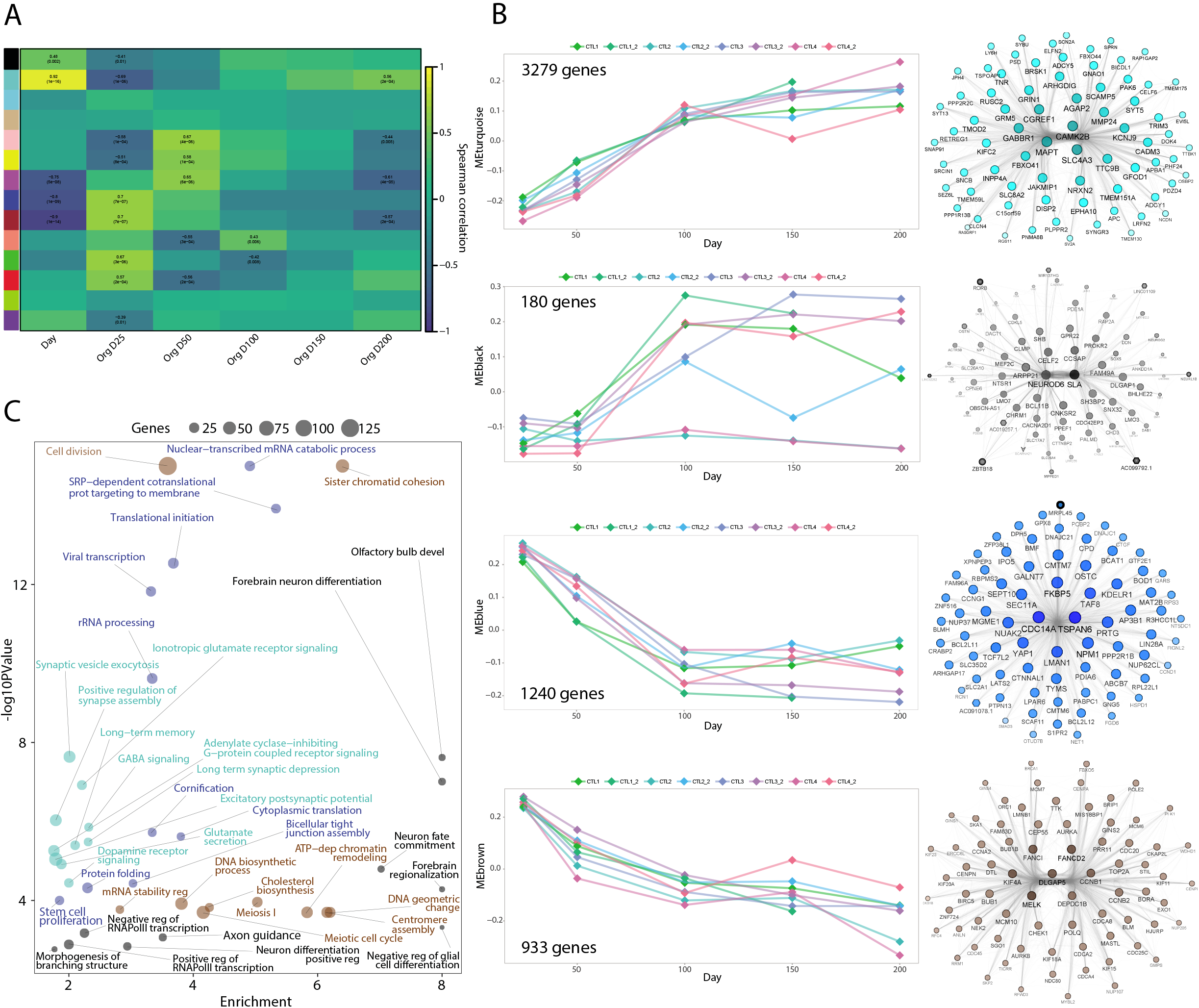
Cortical organoids evolve towards the glutamatergic fate. (**A**) Heatmap showing the correlation between each gene module (summarized as ME) and CBO differentiation stage either as a continuous variable (Day) or a categorical variable for each time-point. Correlation coefficient and p-values are reported for significant correlations (Spearman p-value < 0.01). (**B**) Characterization of CBO_Turquoise, CBO_Black, CBO_Blue and CBO_Brown gene modules. For each module, the ribbon plot visualizes the behaviour of the module eigengene throughout differentiation time-points; each dot represents a data point, while the line connects timepoints of the same replicate (line and differentiation batch). Network reconstruction for the top-75 genes of each module, selected according to the intramodular connectivity value. Degree, closeness, betweenness and eigenvector are represented by node label transparency, node colour darkness, node border width, and node size/node label font size, respectively. Node shape represents gene biotype, with protein coding genes as circles and non-coding genes as squares. (**C**) Bubble plot depicting the results of GO enrichment analysis for the four modules: p-values and enrichment scores are reported.

GO enrichment analysis for CBO_Turquoise and CBO_Black retrieved terms related to neuronal fate commitment and maturation. Network reconstruction confirmed CBO_Turquoise hub genes as related to neurotransmission and synaptic function (e.g. NRXN2, AGAP2, SLC4AE, GABBR1 and GRM5), while the CBO_Black network comprised several transcription factors related to excitatory neuron identity (NEUROD6, SLA, BCL11B, RORB) (**Figure 3B-C**).

CBO_Brown and CBO_Blue ME were instead decreasing along differentiation; the first was associated to DNA replication and cell cycle, while the second to more general functions such as transcriptional and translational regulation (**Figure 3B-C**).

WGCNA also detected modules characterized by non-monotonic trends through differentiation. Among them, CBO_Green and CBO_Red showed levels dropping from Day25 to Day50, and increasing again at late stages. Both modules were enriched in genes related to cell adhesion and ECM organization (**Supplementary Figure 3 B-C**).

In summary, we time-resolved the transcriptional evolution of CBO by identifying co-expression patterns and transcriptional hubs driving their differentiation. We found overall consistency among lines from independent individuals and replicates. Networks, their behaviour, and relevance of every gene composing them are available in the **Supplementary File 2**.

### Benchmarking of brain organoids against prenatal human corticogenesis reveals heterochronicity of differentiation across protocols

To compare CBO against other protocols and evaluate them versus the fetal cortex, we selected three external BO datasets for which RNAseq was publicly available. These datasets, different for degree of patterning and culture conditions, included samples at comparable time-points (from 0 to 100 days). The selected BO datasets were: i) minimally-guided neural organoids (MGO)^12^; ii) forebrain organoids (FO)^18^; iii) telencephalic aggregates (TA)^19^, including a first step of 2D differentiation (**Figure 4A and Supplementary Figure 4A-D)**.

**Figure 4:**
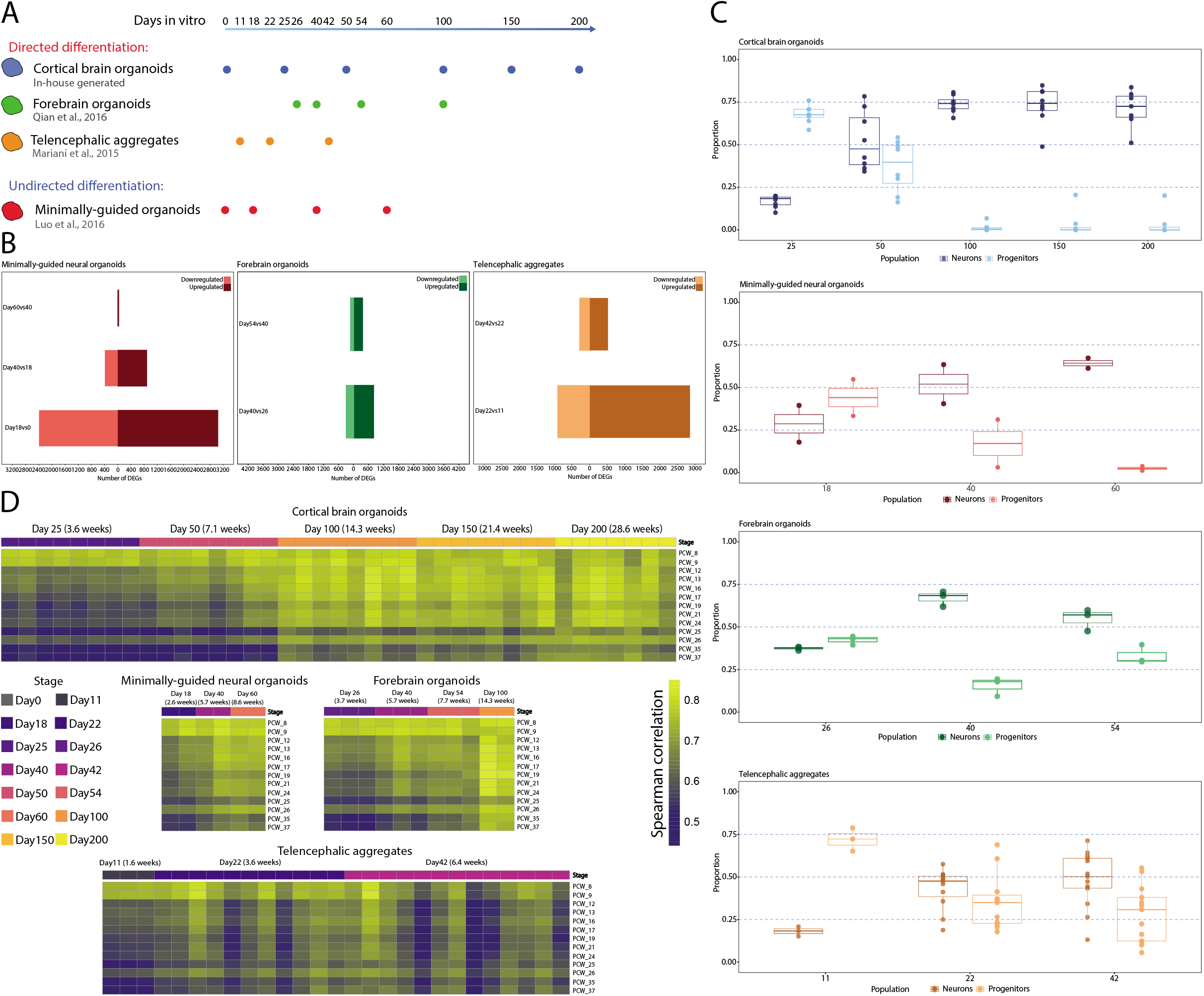
BO differentially recapitulated the timing of corticogenesis. **(A)** Schematic representation of in-house and external BO datasets and relative differentiation timepoints (CBO: Cortical Brain Organoids; FO: forebrain organoids; TA: telencephalic aggregates; MGO: minimally-guided neural organoids). **(B)** Stage-wise differential expression analysis in MGO, FO and TA. Barplots reporting the number of DEGs (FDR < 0.05, log2FC > 1 as absolute value), shown as up-regulated and down-regulated genes for each comparison. **(C)** Boxplots reporting the fraction of Progenitors and Neurons estimated by bulk deconvolution in each organoid dataset. **(D)** Correlation analysis of organoid datasets against the BS fetal cortex. Heatmaps visualize the correlation coefficient (Spearman correlation) for each organoid data point and each BS PCW.

Stage-wise DEA revealed also for MGO, FO, and TA a decrease in the number of DEGs as differentiation progresses (**Figure 4B**). However, the slow-evolving phase, which started between day 100-150 in CBO (**Figure 2C**), was anticipated for all other protocols (day 40-60) (**Figure 4B**). Bulk deconvolution focusing on two broad cell populations (early progenitors and neurons) showed progenitors as predominant in all models at early stages (**Figure 4C**). In CBO, the proportions of the two populations were comparable at day 50, with neurons prevalent from day 100 onwards. MGO, FO and TA showed a similar but accelerated dynamic compared to CBO, suggesting a more rapid transcriptional maturation.

We then tested the transcriptional similarity of each model towards BS fetal cortex (**Figure 4D**). Whole-transcriptome correlation showed for CBO a gradual increase of similarity towards mid and late PCW over time. We observed a more time-compressed evolution among MGO and FO, which already by day 60 showed an extent of similarity with late PCW that CBO reached only by day 100. For CBO and TA, the dataset encompassed organoids generated from different control individuals. CBO demonstrated robust reproducibility across genetic backgrounds and batches of differentiation, while TA were less tolerant to interindividual variability. These observations were further confirmed by using CBO as reference (**Supplementary Figure 4E**).

Overall, several lines of evidence pointed towards heterochronicity across BO models in recapitulating the transcriptional modulations of corticogenesis.

### Gene signatures specific for brain cell subpopulations and functions unveil dynamics of brain organoid differentiation

We then examined how specific hallmarks of brain cell sub populations and functions evolve during BO differentiation. To this end, we compiled a catalogue of genes by manually curating new signatures from literature (**Figure 5**) and by using signatures proposed in published studies^9,13^. We observed their expression dynamics across BO and the developing cortex. Expression levels of signature genes are reported in **Figure 5** and **Supplementary Figure 5**; the dynamics characterizing each model compared to the fetal cortex are detailed in **Supplementary Table 2**.

**Figure 5:**
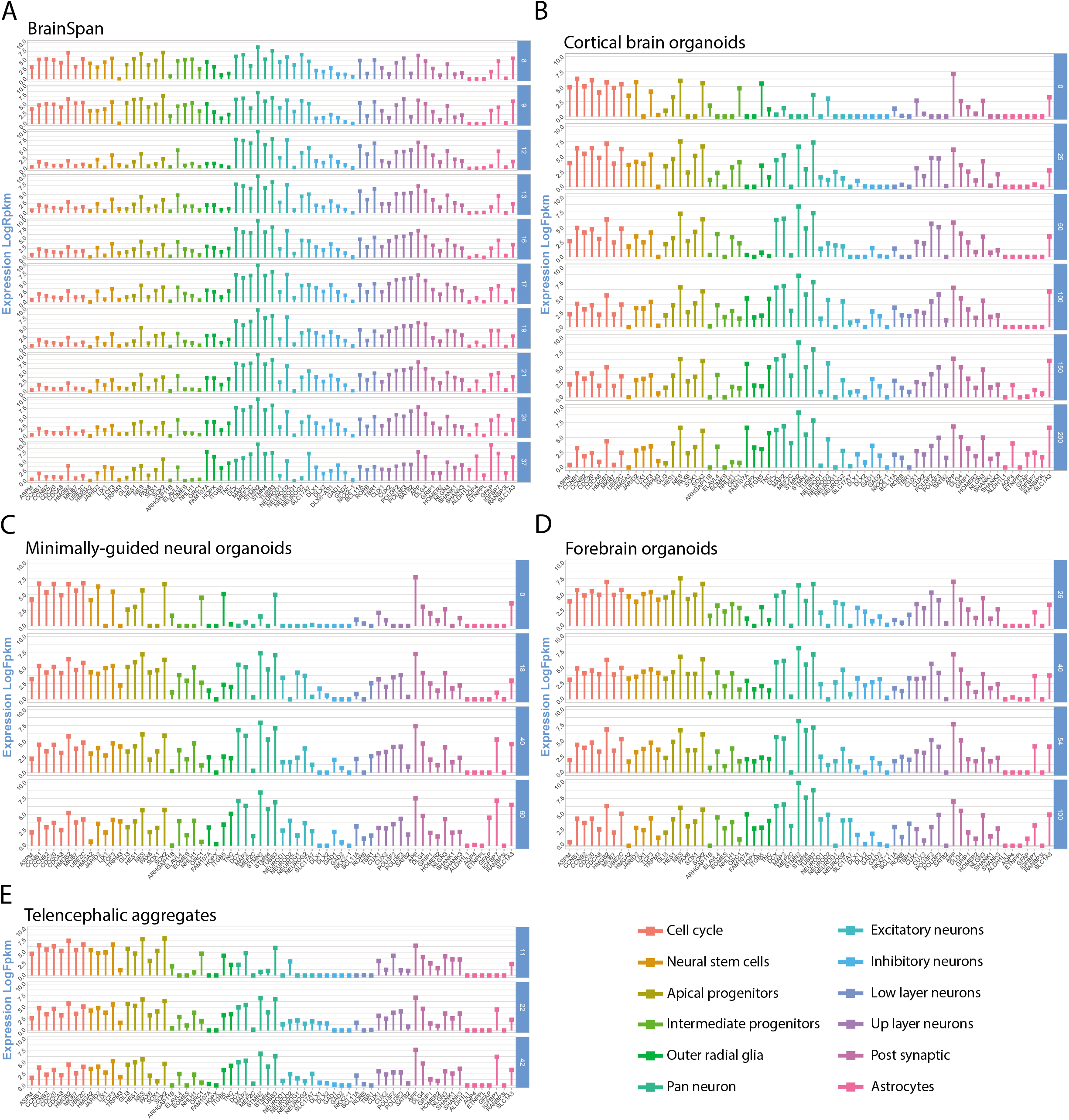
Gene expression levels of signatures related to cell population identity and function in BO. Expression levels (Log2Fpkm or Log2Rpm) in BS **(A)**, CBO **(B)**, MGO **(C)**, FO **(D)** and TA **(E)** along time-points are reported in each dataset as the mean value across replicates. Each bar colour corresponds to a specific signature, as reported in the plot legend.

Fetal cortex showed drastic reduction of cell cycle, neural stem cell, apical progenitor, and intermediate progenitor gene levels between PCW 9-12, in line with WGCNA results. We found in BO models a less abrupt reduction of cell cycle and apical progenitor markers. The intermediate progenitor signature was found to peak at different time-points across models.

Excitatory neuron markers were particularly expressed at early PCW in the fetal cortex, a trend mirrored by CBO; MGO, FO and TA showed a more persistent expression throughout differentiation (**Supplementary Table 2**).

oRG genes were well expressed in BS at PCW 8-9, decreased from PCW 12 and increased again at later stages, when astrocyte markers also appeared. CBO showed constant up-regulation of the whole oRG signature starting from day 100, while for FO this was observed already at day 40; MGO and TA showed a partial expression at the available time-points. Astrocyte markers were inconsistently detected in BO, except for CBO that showed stable expression at day 150 and 200.

Considering that the examined BO protocols were based on different degrees of patterning and culturing conditions, we then looked at brain area specific genes, markers of off-target tissues, markers of neurotransmission, and stress-related genes (detailed in **Supplementary Figure 5, Supplementary Figure 6, Supplementary Table 2**). As markers of metabolic stress, we looked at the expression of glycolytic and ER-stress genes^13^. We found an overall stable expression of those genes throughout cortical development as well as in BO, without hints of strong up-regulation at advanced time-points.

### Brain organoids differentially capture distinct transcriptional patterns of fetal corticogenesis

The knowledge base of gene co-expression modules we reconstructed for corticogenesis and CBO differentiation allowed us to directly compare modules identified in cortex/CBO and follow their behavior in the other BO models. First, we measured the overlap between the two sets of WGCNA modules (**Supplementary Figure 7)**. Fetal BS_Turquoise, BS_Pink, BS_Midnight-blue, and BS_Grey60 genes, steadily increasing their expression along cortigogenesis, significantly overlapped with organoid CBO_Turquoise, which showed the same trend. Similarly, BS_Yellow and BS_Black, functionally associated to cell cycle, showed steady decrease over time and overlapped with CBO_Brown, which had similar behaviour and functional characterization (**Supplementary Figure 7**).

We then analyzed the behaviour of these BS modules in each organoid dataset by following their ME across differentiation, revealing a close resemblance to fetal cortex trends (**Figure 6A-B**). In a complementary approach, we applied the same analysis on the most relevant modules detected in CBO. The behaviour of CBO_Turquoise and CBO_Brown in fetal cortex and other BO protocols was consistent with the one described in CBO. CBO_Black genes induction was variable across models and replicates (**Supplementary Figure 8A**). CBO_Blue genes behaved similarly in all BO, and showed instead a non-monotonic behaviour in BS, with peaks of expression at very early and very late PCW (**Supplementary Figure 8B**).

**Figure 6:**
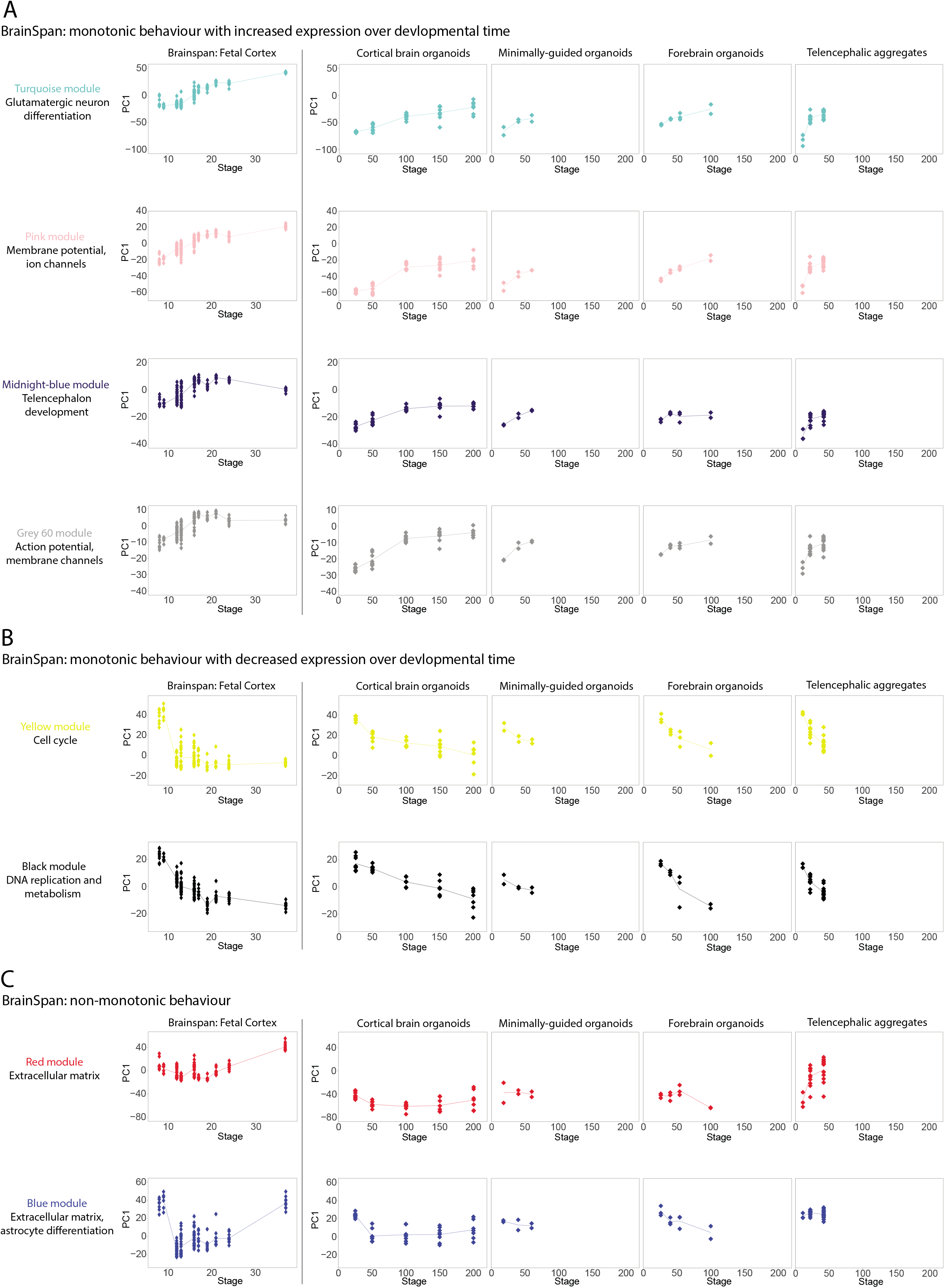
Behaviour of the module eigengene for BrainSpan gene co-expression patterns in BO. **(A)** Visualization of BS_Turquoise, BS_Pink, BS_Midnightblue and BS_Grey60 in prenatal fetal cortex as well as in the BO datasets. PCA was performed to calculate the module eigengene on the BrainSpan and then applied to each BO dataset. Each dot represents a data point, while the line connects the median value for each post-conceptional week (BS) or differentiation day (BO). The same analysis and visualization was applied to BS modules with a decreased expression along development **(B)** or with a non-monotonic behaviour **(C)**.

Finally, we analyzed how BO recapitulated patterns of non-monotonic expression through time. BS_Red, BS_Blue and CBO_Red, CBO_Green significantly overlapped between the two networks. The visualisation of BS_Red and BS_Blue trends in BO revealed differences across protocols. CBO mostly displayed the same U-shape found in BS, although with a milder drop at intermediate stages. Conversely, MGO showed very little variation, FO showed a drop at day 100, and TA showed an opposite trend for the red module and no change for the blue one (**Figure 6C**). Likewise, CBO and BS behaved similarly for the set of CBO_Red and CBO_Green genes, while these modules showed no, little or opposite variation in MGO, FO and TA (**Supplementary Figure 8C**). Among the functions of this set of modules, ECM and cell adhesion were prevalent.

Together, these results suggested that monotonic gene expression dynamics related to neuron specification and progenitor proliferation are shared between fetal cortex and CBO and are largely recapitulated by the analyzed BO paradigms. Conversely, trends of non-monotonic expression are largely recapitulated in CBO and less robustly in other BO systems at the examined time-points.

## DISCUSSION

Despite the complexity of neuropsychiatric conditions, in terms of both phenotypic and time course heterogeneity, it is now well established that BO can allow the discovery of relevant developmental endophenotypes that are starting to illuminate disease pathogenesis with first inroads also in terms of patients’ stratification and drug discovery^8,40^. The ambition is thus rapidly progressing from the initial focus on monogenic syndromes of high penetrance to the more high throughput and challenging interrogation of polygenic loadings for mental health vulnerability, including in terms of the developmental antecedents of its later onset manifestations. Yet, the more central neurodevelopment becomes to our understanding of mental health, the more we need to gain a full understanding of its experimental recapitulation *in vitro*, so as to define clear benchmarks across systems and thereby empirically guide the most appropriate disease modelling designs. Towards this goal, here we first defined co-expression patterns characterizing human corticogenesis *in vivo* and *in vitro* and then organized them in interactive tables where gene relevance can be visualized, together with the overall behaviour of these patterns along fetal cortex development and CBO differentiation (**Supplementary File 1 and 2**). The generation of this knowledge base allowed us to perform cross-comparisons between BS, CBO and other BO paradigms, quantifying the preservation of co-expression patterns of corticogenesis in these models. To our knowledge, this represents the first resource of such kind. Similar analyses were performed on BS and BO^10,12^, however without focus on prenatal cortex, accessibility to gene modules composition, or prioritization of transcriptional hubs (both known and novel). Thus, this knowledge base templates an open framework for the benchmarking of modelling studies and serves as an easy access platform to interrogate when choosing the experimental system that best suits specific modeling needs or research questions. Several studies provided insights on the variability of BO, with considerable efforts of standardization. Our results demonstrated CBO reproducibility in recapitulating the transcriptional landscape of prenatal corticogenesis, in line with other reports^10,41^, with a developmentally relevant timing. We observed that TA were more variable compared to CBO in terms of cell composition and global transcriptome when considering hiPSCs from different individuals. The same evaluation could not be carried out for MGO and FO due to the availability of a single genetic background.

Several analytical approaches (DEA, whole-transcriptome correlation with fetal cortex, bulk deconvolution) revealed a fast-evolving phase followed by a slow-evolving one during CBO differentiation. Among genes showing modulation also in the slow-evolving phase, we found up-regulation of GABAergic neuron markers (such as CALB2, SCGN, SLC6A11), in line with previous studies^15,36,37,42^. Among the functional categories emerging when analysing the slow-evolving phase, we observed an up-regulation of ECM-related genes. We also observed upregulation of genes related to mesoderm, however the majority of them is expressed also in astrocytes/reactive astrocytes^32,43,44^. The developmental timing of astrocytes appearance in CBO (day 100-150 - week 14.3-21.4) aligned with the human prenatal cortex. The interplay between neuronal and astrocytic cells is increasingly recognized in neuropsychiatric disorders^45–47^, therefore the characterization of their differentiation in BO compared to fetal cortex is important to guide disease modelling.

In the limited time-frame available for external BO datasets, we confirmed their biphasic evolution. However, we observed heterochronicity among protocols. We compared similar *in vitro* ages for all protocols analyzed, with the CBO dataset including also very late stages, and observed that MGO, FO and TA reached the slow-evolving phase earlier than CBO, with a greater proportion of neurons at earlier time-points. Additionally, MGO and FO globally correlated earlier with late stages of corticogenesis and CBO differentiation.

A possible explanation for these findings could be related to the maintenance of a more immature stage for longer time due to the prolonged use of EGF/bFGF during CBO differentiation, which however ended up matching more faithfully the *in vivo* counterpart. It is also known that iPSCs can have different propension to neuronal induction depending on the line itself and on culture conditions^48,49^. However, both CBO and TA were generated from multiple hiPSCs lines reprogrammed from different individuals, thereby making improbable that differences across cell lines represent the main source of the observed heterochronicity among protocols. To our knowledge, this is the first description of such temporal dynamics in BO. Other reports comparing different paradigms focused on more heterogeneous time-points across protocols and other aspects of recapitulation^13–15^. Temporal dynamics recapitulation has remarkable relevance for disease-modelling, given the emergent convergence of molecular phenotypes of delay or acceleration in neuronal differentiation characterizing different clinical conditions^50,4^, and the eventuality of masking disease-relevant phenotypes if they are not correctly preserved^51^. Moreover, the tight link between neurodevelopment and risk/resilience of neuropsychiatric disorders positions the *in vitro* recapitulation of developmental timing at the core of the choice between BO systems, also with a view to balance the preservation of accurate developmental timing vis a vis the accelerated yield of more mature cell types involved in later onset neuropsychiatric disorders. Our analyses started to elucidate these dynamics in BO, suggesting that CBO differentiation aligns with the *in vivo* temporality of corticogenesis.

Cellular stress in BO was previously reported by scRNAseq with the up-regulation of glycolytic and endoplasmic reticulum stress signatures in several paradigms compared to primary samples^13,14^. However, we detected a stable expression of stress-related signatures throughout differentiation in CBO, in all other BO datasets corroborating the hypothesis of a homeostatic metabolic state in organoids rather than a deleterious stress increase over-time, in line with a recent report^10^.

Cross-visualization of monotonic BS co-expression patterns in CBO and *vice versa* revealed concordance in their behaviour, a finding observed also for the other BO datasets. Conversely, modules with more complex temporal patterns were better recapitulated in CBO. Functional characterization of these modules pointed towards astrogenesis and ECM as predominant terms.

We also processed primary fetal tissues in-house, allowing direct comparisons against CBO because of the identical processing of the two datasets. The visualization of cortex-specific modulated genes in CBO highlighted again ECM. ECM genes have dynamic expression during corticogenesis, and have a role in regulating cortical folding, neuronal progenitor proliferation, and neuronal migration^52,54^. They have high expression in germinal zones and maturing/mature astrocytes^52–55^ and are important for the evolution of the human brain, with progenitor cells expressing ECM components at higher levels than in mice^56^. ECM also plays critical roles in synaptic plasticity and electrophysiological properties^57,58^, and its alteration has been linked to neuropsychiatric disorders^59,60^. Our exposure of the differential regulation of ECM genes in BO compared to fetal cortex provides therefore a new relevant insight to consider when interpreting neuropsychiatric disease modelling datasets.

Taken together, our results contribute to the definition of transcriptional footprints and dynamics specific to prenatal corticogenesis, representing a collection of prioritized known and novel hubs categorized in well-defined functional domains and made available as interactive tables (**Supplementary file 1 and 2**). Our approach describes the extent of *in vivo/in vitro* alignment of developmentally relevant processes and temporality, highlighting commonalities and diversities of different BO paradigms and providing a resource available for consultation when modelling physiological or pathological human corticogenesis.

## METHODS

### hiPSCs reprogramming

Skin fibroblasts from four healthy individuals were reprogrammed using non integrating self-replicating mRNAs as previously described^61^ (Stemgent, 00-0071) (CTL1, CTL3 and CTL4) or Sendai virus (CTL2) (CytoTune-iPS 2.0 Sendai Reprogramming Kit; Thermo Fisher Scientific, A16517). Fibroblasts of CTL1 (phenotypically normal male without intellectual disability or other physician-diagnosed neuropsychiatric diagnosis) were received from BC Children’s Hospital, Vancouver. CTL2 (male) hiPSC line was received from the Wellcome Trust Sanger Institute. CTL3 (male) hiPSC line was received from the university of Sheffield. Fibroblasts of CTL04 (female) were received from Genomic and Genetic Disorders Biobank at the IRCCS Casa Sollievo della Sofferenza, San Giovanni Rotondo (Italy, affiliated to the Telethon Network of Genetic Biobanks). hiPSC were cultured in TeSR-E8 medium (Stemcell technologies, 05990), with daily media change, at 37 °C, 5 % CO2 and 3 % O2 in standard incubators. hiPSC were grown on matrigel-coated dishes (Corning, 354248) and passaged using ReLeSR™ (Stemcell technologies, 05872).

### CBO differentiation

CBO were generated using an adaptation of the previously described protocol published by Pasca et al in 2015^17^, introducing orbital shaking on day 12 of differentiation as described in^16^. Briefly, hiPSC were grown on feeders for 3-4 days in a medium composed of 80% DMEM/F12 medium (1:1), 20% Knockout serum (Thermo Fisher Scientific, 10828028), 1% Non-essential amino acids (NEAA, Lonza BE13-14E), 0.1 mM cell culture grade 2-mercaptoethanol solution (Thermo Fisher Scientific, 31350010), 2 mM L-Glutamine (Thermo Fisher Scientific, 25030081), P/S 100 U/mL, and FGF2 at a final concentration of 20 ng/mL (Thermo Fisher Scientific, PHG0021). Daily media change was performed. Embryoid bodies (EB) were generated by detaching hiPSC with dispase for 40 minutes and plating on ultralow attachment 10 cm plates (Corning, 3262) in the first differentiating medium composed of 80% DMEM/F12 medium (1:1), 20% Knockout serum, 1% NEAA, 0.1 mM cell culture grade 2-mercaptoethanol solution, 2 mM L-Glutamine, P/S, 100 U/mL, 7,5 μM Dorsomorphin (MedChem express, HY-13418A), 10 μM TGFβ inhibitor SB431542 (MedChem express, HY-10431), and ROCK inhibitor 5 μM. EB were grown in normal oxygen incubators. EB were left undisturbed for 1 day and at 48h media change was performed with differentiation medium 1 without ROCK inhibitor. Dorsomorphin and TGFβ inhibitor are used to perform Dual-SMAD inhibition, pushing neuroectoderm specification. Dual-SMAD inhibition was performed for a total of 5 days, with daily media change. On day 6 the second differentiation medium was added until day 25 with daily media change for the first 12 days, and then every other day. The second differentiation medium was composed of neurobasal medium (Thermo Fisher Scientific, 12348017) supplemented with 1X B-27 supplement without vitamin A (Thermo Fisher Scientific 12587001), 2 mM L-Glutamine, P/S, 100 U/mL, 20 ng/mL FGF2and 20 ng/mL EGF (Thermo Fisher Scientific, PHG0313). Human FGF2 and EGF were used to amplify the pool of neural progenitors. On day 12, CBO were moved to ultra-low attachment 10 cm dishes and grown on shakers to enhance oxygen and nutrient supply. On day 26, FGF2 and EGF were replaced with 20 ng/mL brain-derived neurotrophic factor (BDNF, Peprotech 450-02) and 20 ng/mLneurotrophin-3 (NT3, Peprotech 450-03) to promote differentiation of neural progenitors towards the glutamatergic fate. From day 43 onwards, BDNF and NT3 were removed and from day 50 the medium was supplemented with Amphotericinβ to prevent mold formation.

### Culture conditions for fetal cortical cell

Human fetal neural stem cell cultures were derived and maintained as previously described^62^. They were derived from the cortex of WGA 11 and 19, male embryos. They were cultured in the following medium DMEM/F12 medium (1:1), P/S (100 U/mL), 0.1 mM cell culture grade 2-βmercaptoethanol solution, 1% NEAA, 0.5% N2 supplement, 1X B27 Supplement 100X (Thermo Fisher Scientific, 17504-044), 0,012% Bovine Albumin Fraction V (Thermo Fisher Scientific, 15260-037), 1,5 g/L glucose (Sigma-Aldrich, G8644). Washes for this type of cells were performed with a medium composed of DMEM/F12 medium (1:1), P/S (100 U/mL) and 0,015% Bovine Albumin Fraction V.

### Download of external datasets

<u>BrainSpan dataset</u>: pre-processed Rpkm values for the BrainSpan Atlas were downloaded from here: http://www.brainspan.org/static/download.html. Dataset organisation was described in the following white paper: https://help.brain-map.org/display/devhumanbrain/Documentation, Developmental Transcriptome.

<u>CO, FO and TA datasets</u>: bulk RNA sequencing data were downloaded from Gene Expression Omnibus using the relative article accession numbers (GSE82022, GSE80073, GSE61476, respectively).

### RNA extraction and library preparation for RNA-seq

Total RNA was extracted from snap-frozen pellets of CBO at day 25, 50, 100, 150, 200, fetal cortical progenitors and fetal brain tissues using the RNeasy Mini Kit (Qiagen, 74104). Purified RNA was quantified using a NanoDrop spectrophotometer and RNA quality was checked with an Agilent 2100 Bioanalyzer using the RNA nano kit (Agilent, 5067-1512). Library preparation for RNA sequencing was performed according to TruSeq Total RNA sample preparation protocol (Illumina, RS-122-2202), starting from 250 ng - 1 μg of total RNA. cDNA library quality was assessed at on Agilent 2100 Bioanalyzer, using the high sensitivity DNA kit (Agilent 5067-4626). Libraries were sequenced with the Illumina Novaseq machine at a read length of 50 bp paired-end and a coverage of 35 million of reads per sample.

### RNAseq quantification for CBO, in-house fetal dataset, external brain organoids datasets

RNAseq FASTQ data were quantified at the gene level using Salmon (version 0.8.2,^63^). GRCh38 Genecode 27 was used as reference for quantification and annotation. Data will be available in ArrayExpress public repository upon publication.

### BrainSpan correlation analysis across developmental stages

The analysis was applied on BS specimens from prenatal and postnatal cortex. After selecting protein-coding genes, the mean expression was calculated for each stage (taking into account all the samples and sub-areas). Spearman correlation across samples was calculated and the correlation coefficient was visualized by heatmaps using pheatmap R package (https://CRAN.R-project.org/package=pheatmap).

### Principal Component Analysis

<u>BrainSpan</u>: after selecting only samples from prenatal cortex, a filtering to discard not-expressed or low-expressed genes was applied by keeping genes with expression of at least 1 Rpkm in at least 1/4 of the samples (16824 genes selected).

<u>Internal fetal dataset and CBO</u>: a total of 66 samples considering CBO and internal fetal samples was used to perform PCA. Gene filtering using a threshold of 2 counts per million reads (cpm) in at least 2 samples was used, resulting in 17759 analysed. <u>CBO dataset</u>: a total of 43 samples considering CBO dataset (from day 25 to day 200). Gene filtering using a threshold of 2 cpm in at least 4 samples was used, resulting 16901 genes used for the analysis.

For all datasets, PCA was computed using R prcomp function. For both BS and CBO PCA, gene loadings for PC1 and PC2 were retrieved from this analysis and the top 35 ones with positive and negative scores were visualised as lollipop graphs. The top 300 genes with the highest positive loading and the top 300 with the highest negative loading were selected to perform GO for biological processes using the TopGO package^64^, relying on Fisher test and Weight01 method. p-value < 0.01 and enrichment > 1.75 were used as thresholds to select significantly enriched GO terms.

### WGCNA

#### BrainSpan

##### Weighted Gene Co-expression network generation and module identification

After selecting only samples from prenatal cortex, not-expressed or low-expressed genes were discarded by keeping genes with expression of at least 1 Rpkm in at least 1/4 of the samples (16824 genes selected). Samples from post-conceptional weeks 25, 26 and 35 were identified as outliers by the sample clustering, and therefore excluded from downstream analyses. Starting from this set of 16824 genes measured in 157 specimens, a gene selection strategy was applied by calculating for each gene the coefficient of variation (CV) across the experimental conditions after log-transformation. A 65-percentile threshold was then imposed, thus selecting the 35% of genes showing the highest values of CV (5889 genes). A signed gene co-expression network was generated relying on WGCNA R package (version 1.64.1,^65^). The correlation matrix was calculated by applying a biweight mid-correlation and then transformed into an adjacency matrix by raising it to the power of β = 18. Topological Overlap Measure was calculated from the adjacency matrix and the relative dissimilarity matrix was used as input for average-linkage hierarchical clustering and gene dendrogram generation. Network modules were detected as branches of the dendrogram by using the DynamicTree Cut algorithm (deepSplit=1; minimum cluster size= 50; PAM stage TRUE; cutHeight 0.998,^66^.

##### Module-trait correlation

As phenotypic trait for module-trait correlation, module eigengenes were related to each sample developmental stage (PCW), considered either as a continuous quantitative variable or dichotomized as a series of categorical variable (dummy variables).

##### Module functional analysis

Gene ontology enrichment analysis for the Biological Process domain was performed on the genes belonging to each module of interest, using the list of 5889 genes selected for network generation as custom reference set. Analysis was performed by TopGO as described above. p-value < 0.01 and enrichment > 1.75 were used as thresholds to select significantly enriched GO terms.

##### Sub-network visualization and analysis in Cytoscape

Node and edge information for the selected modules were exported from the adjacency matrix, selecting for each module the top-75 according to the intramodular connectivity and imposing a threshold of 0.2 as minimum edge weight. Nodes and edges for each module were therefore imported in Cytoscape (version 3.8.2) for centrality analysis (CytoNCA,^67^).

#### CBO

The same pipeline used to perform WGCNA on BS was applied also to the CBO dataset. The analysis was performed on a total of 39 samples with in vitro age spanning from day 25 to day 200. Filtering was applied by keeping genes with an expression of at least 2 cpm in at least 7 samples (15663 genes selected). Coefficient of variation was calculated on log-transformed data and was set to 0.5, resulting in a total of 7831 genes considered for the analysis. The soft threshold β was set to 15. The DynamicTree Cut algorithm parameters used for gene module identification were DeepSplit of 1; minClusterSize 30; PAM stage TRUE; cutHeight 0.999, for a total of 14 modules that were then characterised using the same approach described for BS.

For the generation of the bubble plot relative to CBO turquoise, black, blue and brown modules, REVIGO web tool was employed to summarize GO terms^68^.

### Differential Expression Analyses (DEA)

<u>CBO</u>: DEAs were performed for CBO in a stage-wise approach comparing each differentiation stage with the previous time-point using edgeR. Genes with expression levels higher than 2 cpm in at least 4 samples were tested for differential expression (16901 genes). The information about line identity was used as a covariate in the statistical model. DEGs were selected imposing as thresholds FDR < 0.05 and absolute log2FC > 1. For each comparison, the number of DEGs, splitted in up-regulated or down-regulated, was represented by barplots. Functional enrichment analysis for the Biological Process domain of GO was performed by TopGO for every comparison dividing genes in up- and down-regulated. PValue < 0.01 and enrichment > 2 were used as thresholds to select significantly enriched GO terms.

Visualisation of DEA results between the different sequential CBO comparisons was performed using scatter plots visualizing the log2FC of all tested genes; the color-code was set according to FDR values. Gene resulting differentially expressed in both sequential CBO comparisons (e. g. day 50vs25 against day 100vs50) were identified and divided in 4 quadrants. In this way, the behaviour of genes in common between the two DEAs was analysed, thus finding genes up-regulated in both, down-regulated in both or up-regulated in one and down-regulated in the other. The top 10 protein coding genes in terms of absolute fold change for the four type of behaviours were labelled in the plot.

<u>CO, FO and TA datasets</u>: the same stage-wise DEA approach was applied for MGO, FO and TA, analysing each dataset independently Genes with expression levels higher than 2 cpm in at least 2 samples were tested for differential expression by edgeR (CO: 15339; FO: 16585; TA: 16522 genes). Line identity was used as a covariate for the TA dataset, while for CO and FO only one line was available. DEGs were selected and visualized as described for CBO.

### Cortex-specific genes determination and visualization of their behaviour in CBO

DEAs of the different fetal tissues from our internal cohort versus hiPSC were performed for all areas including at least two samples using edgeR. Genes with expression levels higher than 2 cpm in at least 2 samples were tested for differential expression (17275 genes). DEGs with FDR < 0.05 and absolute log2FC > 3 were considered for further analysis. Cortex-specific DEGs were determined by subtracting to DEGs of the cortex vs hiPSC comparison DEGs found in the analysis between other tissues and hiPSC. Functional enrichment analysis for GO biological processes was performed by TopGO for cortex-specific genes dividing them in up- and down-regulated. PValue < 0.01 and enrichment > 2 were used as thresholds to select significantly enriched GO terms. The behaviour of cortex-specific DEGs in the CBO dataset, including hiPSC, was visualized by heatmap using average-linkage hierarchical clustering for rows.

### Bulk deconvolution

Proportions of cell populations were estimated applying a deconvolution approach based on the SCDC algorithm^69^ using bulk-RNASeq counts as input and a scRNAseq dataset of the developing human cortex^38^ as reference. The single cell raw count expression matrix was filtered to discard low quality cells, by keeping cells compliant with the following threshold: mitochondrial RNA content < 5%; ribosomal protein RNA content < 50%; detected genes >450 and <3000; UMI counts >750 and < 10000. Starting from the clustering annotation performed in the original work, pericytes, microglia, endocytes and oligodendrocyte progenitors were not included, as we did not expect to find these cell type in CBO. The remaining cell populations were then grouped in the following 6 categories: ventricular Radial Glia (vRG); outer Radial Glia (oRG); Intermediate Progenitos (IP); cycling progenitors; interneurons: excitatory neurons. After deconvolution and for visualization purposes, the results were further grouped in two broader categories: progenitors (cycling progenitors + vRG) and neurons (excitatory neurons + inhibitory neurons). Cells with uncertain cell-type assignments were removed (SCDC qc threshold = 0.65). Cell proportions were retrieved using the SCDC_prop function and were visualised as boxplots showing the proportion of every cell type per stage.

### Correlation of BO transcriptome versus BS or versus CBO

Transcriptome-wide correlation was calculated on Fpkm (Rpkm for BS) after selecting protein-coding genes and filtering out lowly-expressed ones (mean expression levels lower than 1 Rpkm in at least 75% of samples (BrainSpan) and 1 Fpkm (Brain Organoids)). Mean expression per gene per stage was calculated for all BS prenatal cortical samples. Correlation was computed using Spearman metrics and visualised as separated heatmaps for each comparison. The same approach was applied to perform correlation of external brain organoid datasets against CBO.

### Literature-curated gene signatures visualisation in BS and BO

Expression values (in log2Fpkm or log2Rpkm) for the gene signatures of interest were retrieved and the mean expression was calculated for each stage of every dataset, then visualized by lollipop plots.

### Module overlap of BS and CBO WGCNA

Genes shared across the CBO and BrainSpan networks were selected for the analysis (2643); overlap across modules of interest was performed. Number of shared genes, odds ratio and p-value across CBO and BrainSpan modules were calculated by the GeneOverlap R package. Results were visualised as dot plot were numbers (shared genes) were shown for odds ratio >1, dots were shown for those having also p-value < 0.05. Color-code was assigned according to OR, dot size varied according to p-value.

### Analysis of BS modules in BO and of CBO modules in BS and other brain BO

Module eigengene for each BrainSpan WGCNA module of interest was calculated in all organoid datasets as a prediction (R function predict) based on the module eigengene of BrainSpan itself. Likewise, module eigengene for each CBO WGCNA module of interest was calculated and predicted in BrainSpan and in external brain organoid datasets. Results were visualised as ribbon plots showing first principal component coefficients for each module along BrainSpan and organoid developmental time-points.

### Statistical analyses

All bioinformatic analyses were performed using R version 3.4.4 except for the bulk deconvolution analysis performed using R version 3.6.1. The statistical details of all analysis can be found in the relative figure legend and text of the result section.

### Data and code availability

Bulk RNAseq data generated in this study have been deposited in ArrayExpress and will be made available upon publication. No new algorithms were developed for this work, which employed well known tools such as edgeR and WGCNA (as documented in the method details). Code snippets for data processing and characterization/visualization of gene modules are already included as supplementary files. Similar code for all other analytical steps was versioned in a GitHub repository that will be made available upon publication (and whose access for reviewing purposes could be granted upon request).

## ACKNOWLEDGEMENTS

The authors wish to thank the Wellcome Trust Sanger Institute, its funders and clinical collaborators, and Life Science Technologies Corporation for providing CTL2 hiPSCs. The authors wish to thank the University of Sheffield for providing CTL3 hiPSCs. The authors wish to thank the Genomic and Genetic Disorders Biobank at the IRCCS Casa Sollievo della Sofferenza, San Giovanni Rotondo for providing CTL4 fibroblasts. This work received funding from the European Research Council (ERC) (DISEASEAVATARS 616441 to GT); ENDpoiNTs, European Union’s Horizon 2020 research and innovation program (grant no. 825759. to G.T.); S.T. performed this work while being PhD student within the European School of Molecular Medicine (SEMM). S.T. was funded by a fellowship FIRC-AIRC three-years fellowship “Assunta Lombardelli Giuliano Mordini” id. 24168. W.T.G. is supported by the British Columbia Children’s Hospital Research Institute by an intramural IGAP research salary award. R.B.B. was supported by a PhD fellowship from the Science Without Borders Program (CAPES, Governo Dilma Rousseff, Brazil). The authors gratefully acknowledge the research participants who donated biopsies and/or tissue samples from themselves and their family members.

## AUTHOR CONTRIBUTIONS

C.C. designed the analytical strategy of transcriptomic data. S.T. designed and performed CBO differentiation. C.C. and S.T. performed bioinformatic analyses. N.C. cultured human fetal primary stem cells and contributed to bioinformatic analyses. E.T., F.T. and N.C. helped with cortical organoids maintenance and library preparation. A.L.T. curated literature-derived gene signatures. W.T.G. provided skin fibroblasts of CTL1, and provided editorial comments on the manuscript. M.G. reprogrammed hiPSCs from CTL1’s fibroblasts. R.B.B. and S.P. provided human fetal primary samples. C.C., S.T., A.L.T., and N.C. contributed to the study design, discussion of the results and critical reading of the manuscript. C.C., S.T and G.T. wrote the manuscript. G.T. conceived, designed, and supervised the study.

## DECLARATION OF INTERESTS

Authors declare no competing interests.

## Notes

### Competing Interest Statement

The authors have declared no competing interest.

