## Supplementary Figures for "Benchmarking brain organoid recapitulation of fetal corticogenesis"



#### KEYWORDS

### Figure S1

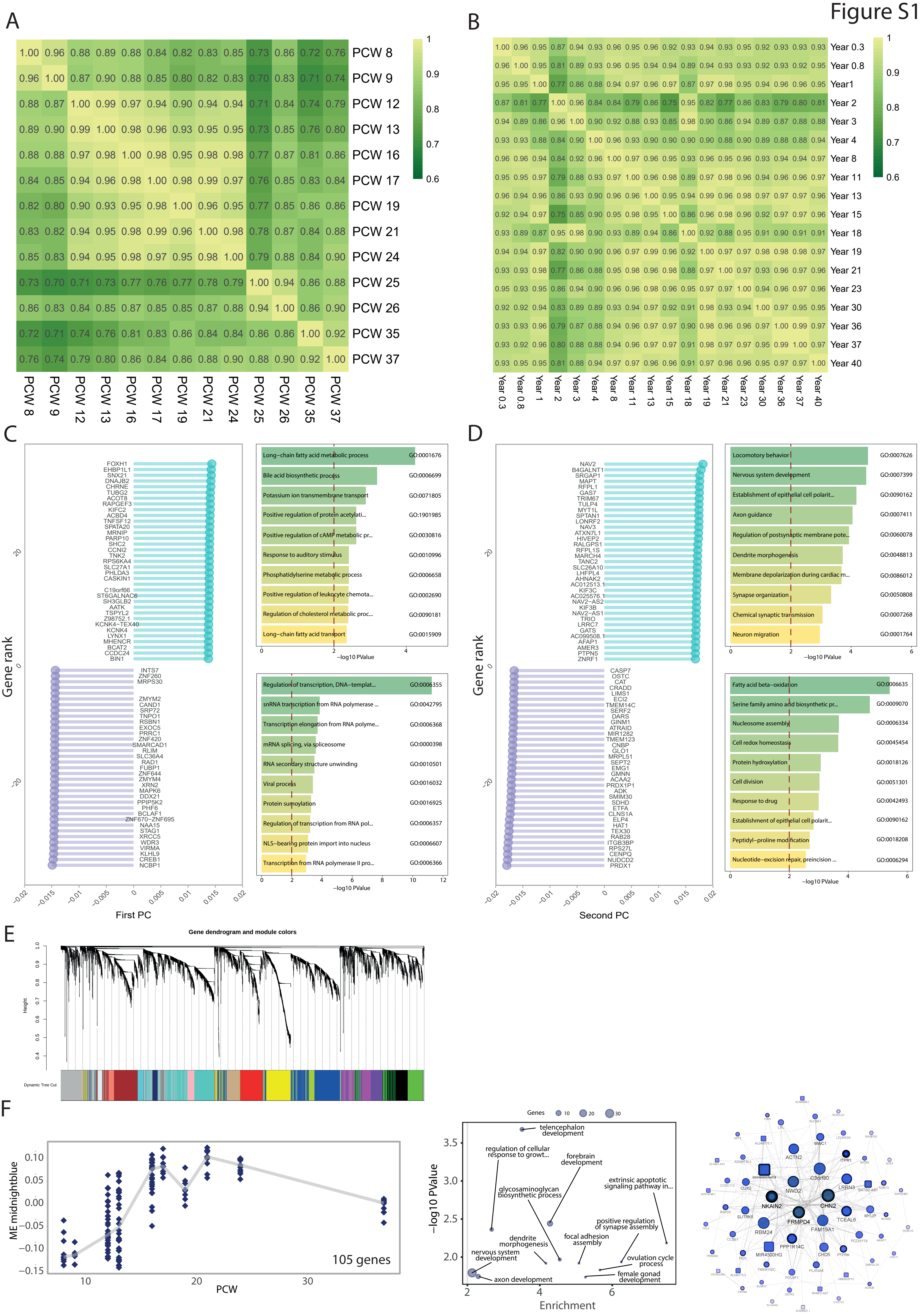

A
PCW 8
PCW 9
PCW 12
PCW 13
PCW 16
PCW 17
PCW 19
PCW 21
PCW 24
PCW 25
PCW 26
PCW 35
PCW 37
B
Year 0.3
Year 0.8
Year1
Year 2
Year 3
Year 4
Year 8
Year 11
Year 13
Year 15
Year 18
Year 19
Year 21
Year 23
Year 30
Year 36
Year 37
Year 40
C
Gene rank
First PC
Long−chain fatty acid metabolic process GO:0001676
Bile acid biosynthetic process GO:0006699
Potassium ion transmembrane transport GO:0071805
Positive regulation of protein acetylati... GO:1901985
Positive regulation of cAMP metabolic pr... GO:0030816
Response to auditory stimulus GO:0010996
Phosphatidylserine metabolic process GO:0006658
Positive regulation of leukocyte chemota... GO:0002690
Regulation of cholesterol metabolic proc... GO:0090181
Long−chain fatty acid transport GO:0015909
−log10 PValue
Regulation of transcription, DNA−templat... GO:0006355
snRNA transcription from RNA polymerase ... GO:0042795
Transcription elongation from RNA polyme... GO:0006368
mRNA splicing, via spliceosome GO:0000398
RNA secondary structure unwinding GO:0010501
Viral process GO:0016032
Protein sumoylation GO:0016925
Regulation of transcription from RNA pol... GO:0006357
NLS−bearing protein import into nucleus GO:0006607
Transcription from RNA polymerase II pro... GO:0006366
−log10 PValue
D
Gene rank
Second PC
Locomotory behavior GO:0007626
Nervous system development GO:0007399
Establishment of epithelial cell polarit... GO:0090162
Axon guidance GO:0007411
Regulation of postsynaptic membrane pote... GO:0060078
Dendrite morphogenesis GO:0048813
Membrane depolarization during cardiac m... GO:0086012
Synapse organization GO:0050808
Chemical synaptic transmission GO:0007268
Neuron migration GO:0001764
−log10 PValue
Fatty acid beta−oxidation GO:0006635
Serine family amino acid biosynthetic pr... GO:0009070
Nucleosome assembly GO:0006334
Cell redox homeostasis GO:0045454
Protein hydroxylation GO:0018126
Cell division GO:0051301
Response to drug GO:0042493
Establishment of epithelial cell polarit... GO:0090162
Peptidyl−proline modification GO:0018208
Nucleotide−excision repair, preincision ... GO:0006294
−log10 PValue
E
Gene dendrogram and module colors
Height
Dynamic Tree Cut
F
ME midnightblue
PCW
105 genes
Genes 10 20 30
−log10 PValue
Enrichment
telencephalon development
regulation of cellular response to growt...
forebrain development
extrinsic apoptotic signaling pathway in...
glycosaminoglycan biosynthetic process
positive regulation of synapse assembly
dendrite morphogenesis
focal adhesion assembly
nervous system development
ovulation cycle process
female gonad development
axon development

#### Supplementary Figure 1

**(A-B)** Correlation analysis of BrainSpan prenatal and postnatal samples. The heatmaps show the correlation coefficient calculated by Spearman correlation across stages, for prenatal (A) and postnatal (B) timepoints, respectively. **(C-D)** Top-35 genes according to positive and negative loading values for the first (C) and second (D) components from PCA analysis. Results of gene ontology enrichment analysis performed for the Biological Process domain on the top genes selected on the basis of their loading value are visualized in the bar plots. **(E)** Gene dendrogram generated on the Topological Overlap Dissimilarity matrix for BS cortex samples; each branch of the dendrogram corresponds to a module of highly interconnected genes, for a total of 17 modules. The row below the dendrogram depicts the module assignment, each identified by a different colour. **(F)** Ribbon plot, GO enrichment analysis and network reconstruction for BS_Midnightblue module.

Figure S2

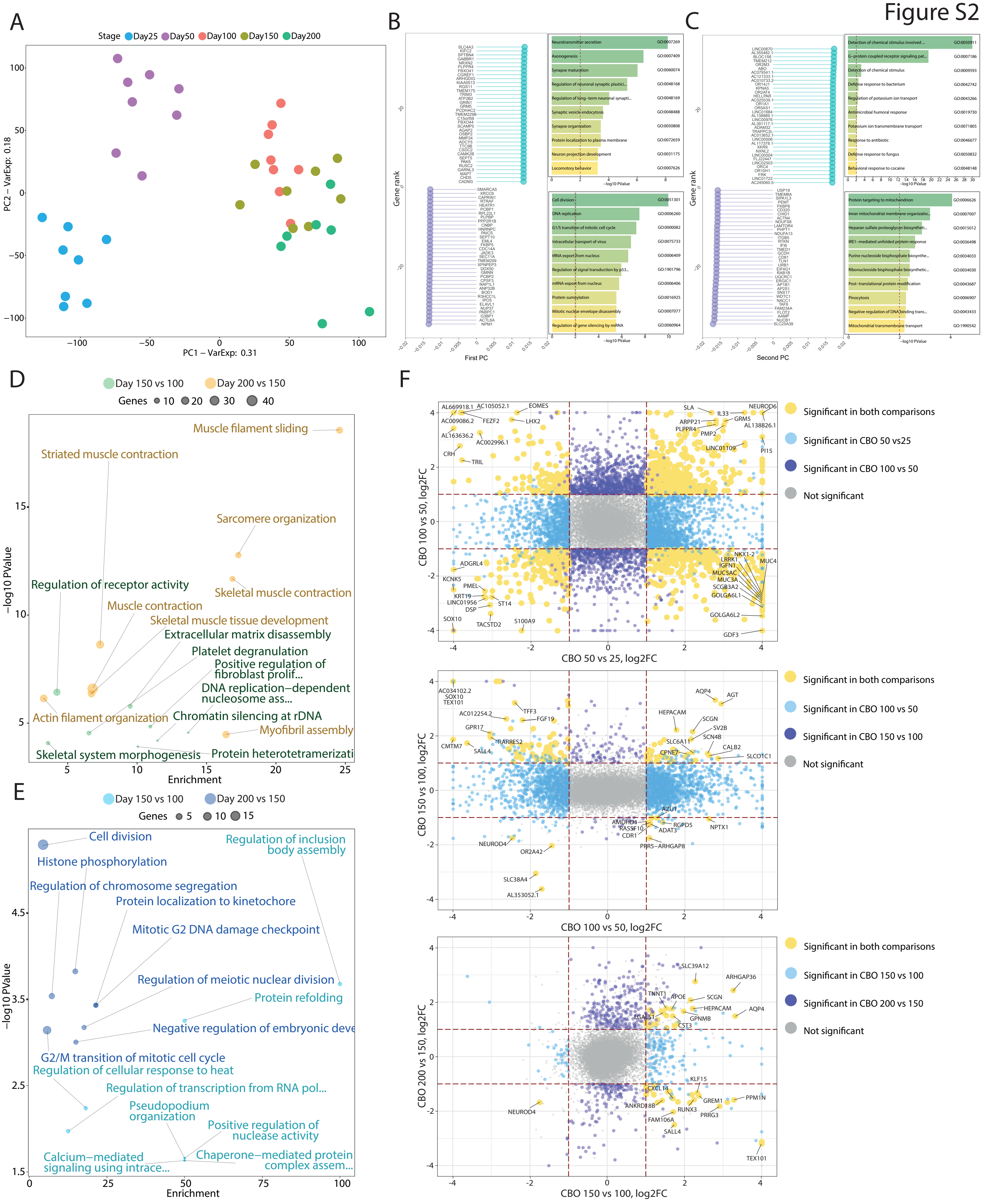
A
Stage Day25 Day50 Day100 Day150 Day200
PC2 – VarExp: 0.18
PC1 – VarExp: 0.31
B
Gene rank
First PC
−log10 PValue
Neurotransmitter secretion
Axonogenesis
Synapse maturation
Regulation of neuronal synaptic plastici...
Regulation of long−term neuronal synapti...
Synaptic vesicle endocytosis
Synapse organization
Protein localization to plasma membrane
Neuron projection development
Locomotory behavior
Cell division
DNA replication
G1/S transition of mitotic cell cycle
Intracellular transport of virus
tRNA export from nucleus
Regulation of signal transduction by p53...
mRNA export from nucleus
Protein sumoylation
Mitotic nuclear envelope disassembly
Regulation of gene silencing by miRNA
C
Gene rank
Second PC
−log10 PValue
Detection of chemical stimulus involved ...
G−protein coupled receptor signaling pat...
Detection of chemical stimulus
Defense response to bacterium
Regulation of potassium ion transport
Antimicrobial humoral response
Potassium ion transmembrane transport
Response to antibiotic
Defense response to fungus
Behavioral response to cocaine
Protein targeting to mitochondrion
Inner mitochondrial membrane organizatio...
Heparan sulfate proteoglycan biosyntheti...
IRE1−mediated unfolded protein response
Purine nucleoside bisphosphate biosynthe...
Ribonucleoside bisphosphate biosynthetic...
Post−translational protein modification
Pinocytosis
Negative regulation of DNA binding trans...
Mitochondrial transmembrane transport
D
Day 150 vs 100
Day 200 vs 150
Genes 10 20 30 40
−log10 PValue
Enrichment
Muscle filament sliding
Striated muscle contraction
Sarcomere organization
Regulation of receptor activity
Skeletal muscle contraction
Muscle contraction
Skeletal muscle tissue development
Extracellular matrix disassembly
Platelet degranulation
Positive regulation of fibroblast prolif...
DNA replication−dependent nucleosome ass...
Actin filament organization
Chromatin silencing at rDNA
Myofibril assembly
Skeletal system morphogenesis
Protein heterotetramerizati
E
Day 150 vs 100
Day 200 vs 150
Genes 5 10 15
−log10 PValue
Enrichment
Cell division
Regulation of inclusion body assembly
Histone phosphorylation
Regulation of chromosome segregation
Protein localization to kinetochore
Mitotic G2 DNA damage checkpoint
Regulation of meiotic nuclear division
Protein refolding
Negative regulation of embryonic deve
G2/M transition of mitotic cell cycle
Regulation of cellular response to heat
Regulation of transcription from RNA pol...
Pseudopodium organization
Positive regulation of nuclease activity
Calcium−mediated signaling using intrace...
Chaperone−mediated protein complex assem...
F
CBO 100 vs 50, log2FC
CBO 50 vs 25, log2FC
Significant in both comparisons
Significant in CBO 50 vs25
Significant in CBO 100 vs 50
Not significant
CBO 150 vs 100, log2FC
CBO 100 vs 50, log2FC
Significant in both comparisons
Significant in CBO 100 vs 50
Significant in CBO 150 vs 100
Not significant
CBO 200 vs 150, log2FC
CBO 150 vs 100, log2FC
Significant in both comparisons
Significant in CBO 150 vs 100
Significant in CBO 200 vs 150
Not significant

#### Supplementary Figure 2

**(A)** PCA on the CBO cohort, with differentiation stage shown as dot color. **(B-C)** Top-35 genes according to positive and negative loading values for the first (B) and second (C) components from PCA analysis and barplots showing functional enrichment results for the Biological Process domain on the top genes. **(D-E)** Functional analysis of up-regulated (D) and down-regulated (E) genes for Day150 vs Day100 and Day200 vs Day150 stage-wise differential expression analysis on CBO. **(F)** Scatterplots representing the relationship of the fold-change across successive time windows. Genes resulting differentially expressed (FDR < 5% and absolute log2FC > 1) in both the two examined comparisons are reported in yellow, while the ones specific for one of the two comparisons are in blue or purple. Gene symbols are reported for the top 10 protein-coding genes for each quadrant according to fold-change.

Figure S3

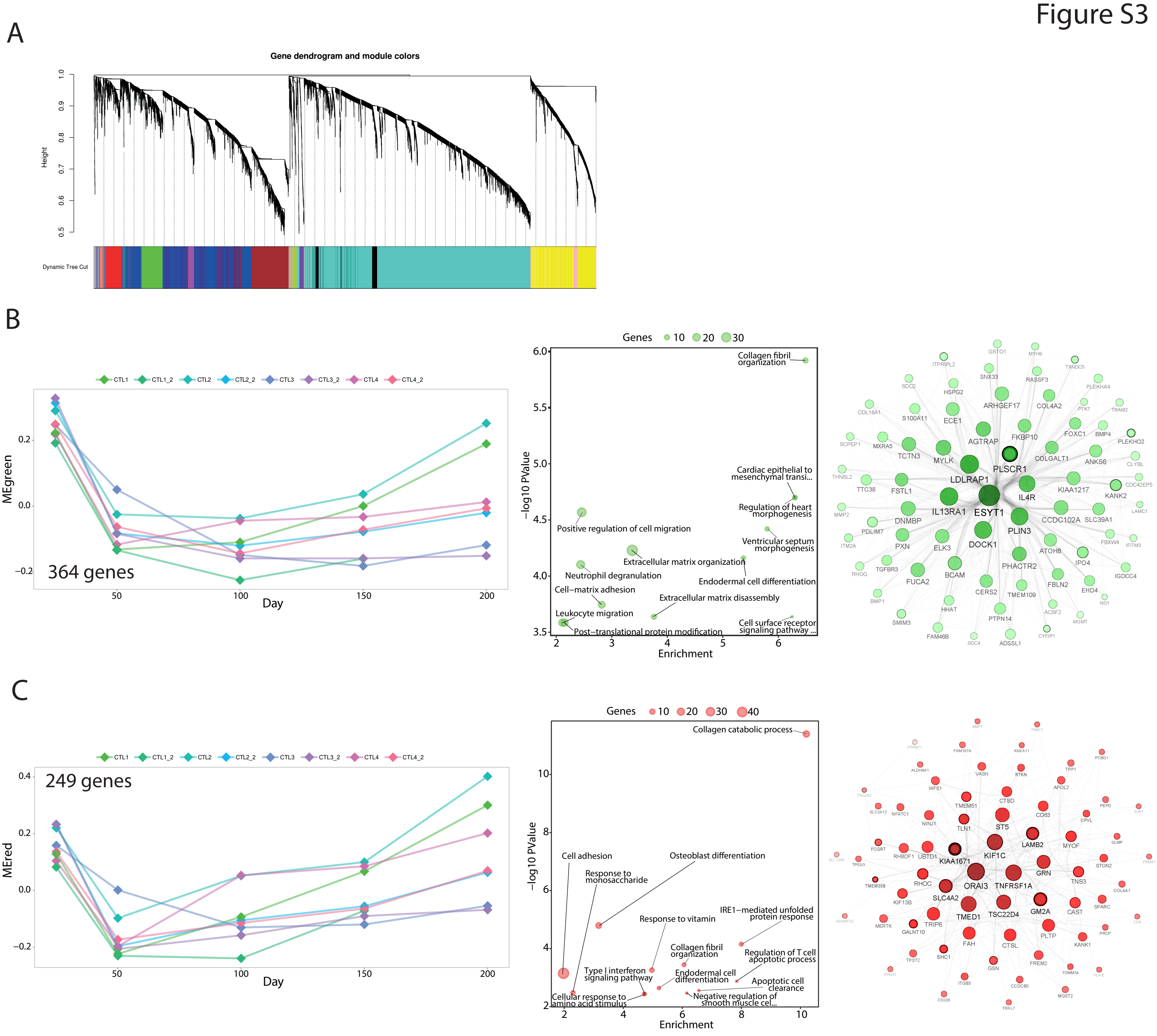

A
Gene dendrogram and module colors
Height
Dynamic Tree Cut
B
CTL1 CTL1_2 CTL2 CTL2_2 CTL3 CTL3_2 CTL4 CTL4_2
MEgreen
364 genes
Day
Genes 10 20 30
−log10 PValue
Enrichment
Collagen fibril organization
Cardiac epithelial to mesenchymal transi...
Regulation of heart morphogenesis
Positive regulation of cell migration
Ventricular septum morphogenesis
Extracellular matrix organization
Neutrophil degranulation
Endodermal cell differentiation
Cell−matrix adhesion
Extracellular matrix disassembly
Leukocyte migration
Cell surface receptor signaling pathway ...
Post−translational protein modification
ESYT1
LDLRAP1
PLSCR1
IL4R
IL13RA1
PLIN3
DOCK1
C
CTL1 CTL1_2 CTL2 CTL2_2 CTL3 CTL3_2 CTL4 CTL4_2
MEred
249 genes
Day
Genes 10 20 30 40
−log10 PValue
Enrichment
Collagen catabolic process
Cell adhesion
Osteoblast differentiation
Response to monosaccharide
Response to vitamin
IRE1−mediated unfolded protein response
Collagen fibril organization
Regulation of T cell apoptotic process
Type I interferon signaling pathway
Endodermal cell differentiation
Apoptotic cell clearance
Cellular response to amino acid stimulus
Negative regulation of smooth muscle cel...
ORAI3
TNFRSF1A
KIF1C
KIAA1671
SLC4A2
TMED1
TSC22D4
GM2A
GRN
LAMB2
ST5

#### Supplementary Figure 3

(**A**) Gene dendrogram generated on the Topological Overlap Dissimilarity matrix for CBO samples; each branch of the dendrogram corresponds to a module of highly interconnected genes, for a total of 14 modules. The row below the dendrogram depicts the module assignment, each identified by a different colour. **(B-C)** Ribbon chart, bubble plot and network reconstruction for the green (B) and red (C) modules.

### Figure S4

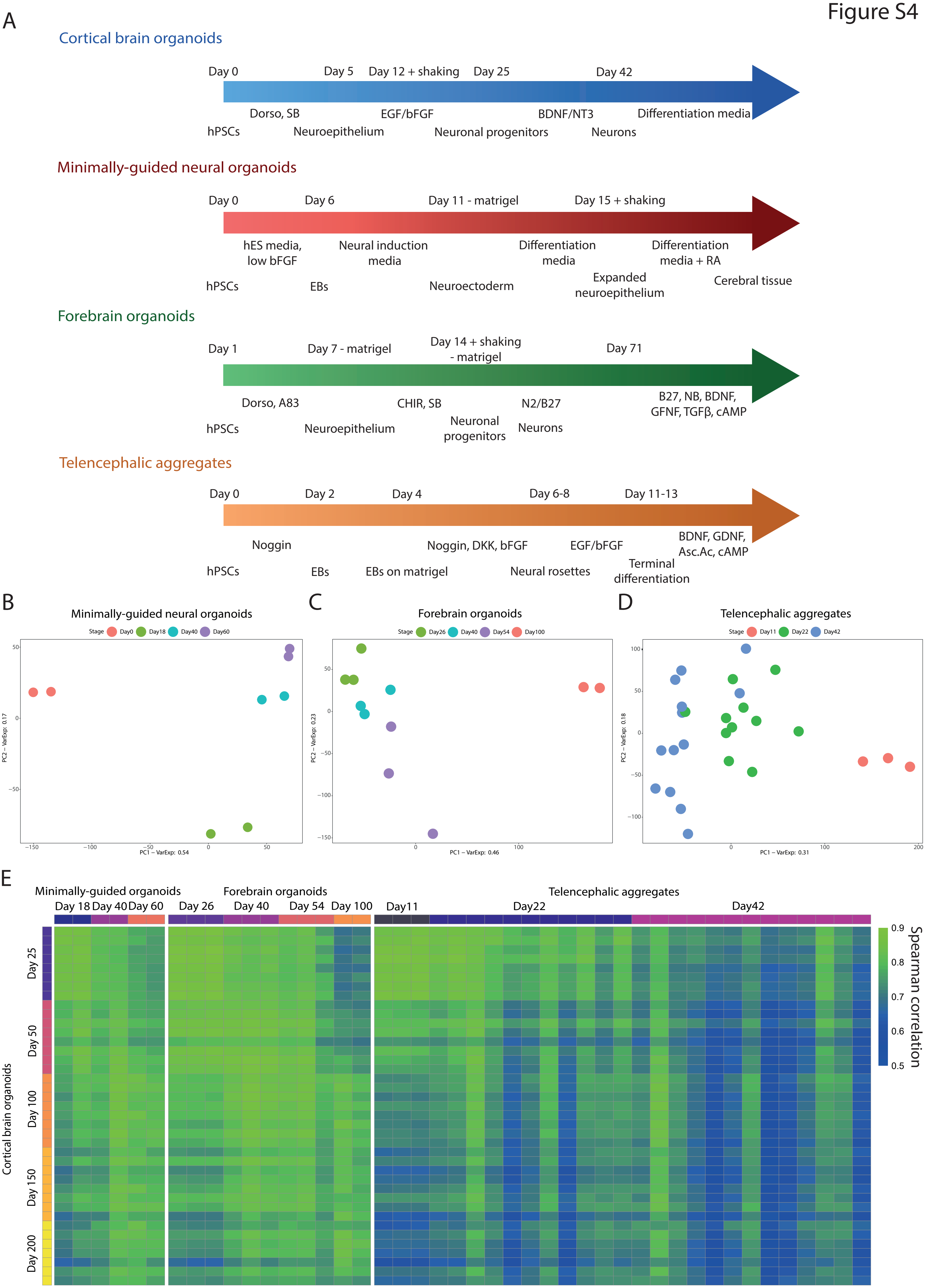
A
Cortical brain organoids
Day 0
Day 5
Day 12 + shaking
Day 25
Day 42
Dorso, SB
EGF/bFGF
BDNF/NT3
Differentiation media
hPSCs
Neuroepithelium
Neuronal progenitors
Neurons
Minimally-guided neural organoids
Day 0
Day 6
Day 11 - matrigel
Day 15 + shaking
hES media, low bFGF
Neural induction media
Differentiation media
Differentiation media + RA
hPSCs
EBs
Neuroectoderm
Expanded neuroepithelium
Cerebral tissue
Forebrain organoids
Day 1
Day 7 - matrigel
Day 14 + shaking - matrigel
Day 71
Dorso, A83
CHIR, SB
N2/B27
B27, NB, BDNF, GFNF, TGFβ, cAMP
hPSCs
Neuroepithelium
Neuronal progenitors
Neurons
Telencephalic aggregates
Day 0
Day 2
Day 4
Day 6-8
Day 11-13
Noggin
Noggin, DKK, bFGF
EGF/bFGF
BDNF, GDNF, Asc.Ac, cAMP
hPSCs
EBs
EBs on matrigel
Neural rosettes
Terminal differentiation
B
Minimally-guided neural organoids
Stage Day0 Day18 Day40 Day60
PC1 – VarExp: 0.54
PC2 – VarExp: 0.17
C
Forebrain organoids
Stage Day26 Day40 Day54 Day100
PC1 – VarExp: 0.46
PC2 – VarExp: 0.23
D
Telencephalic aggregates
Stage Day11 Day22 Day42
PC1 – VarExp: 0.31
PC2 – VarExp: 0.18
E
Minimally-guided organoids
Day 18 Day 40 Day 60
Forebrain organoids
Day 26 Day 40 Day 54 Day 100
Telencephalic aggregates
Day11 Day22 Day42
Cortical brain organoids
Day 25
Day 50
Day 100
Day 150
Day 200
Spearman correlation
0.9
0.8
0.7
0.6
0.5

#### Supplementary Figure 4

**(A)** Schematic representation of the differentiation steps of the four BO protocols analyzed. **(B-D)** PCA of the MGO (B), FO (C) and TA (D) datasets. **(E)** Heatmap depicting the results of the whole-transcriptome correlation analysis between CBO data and external BO protocols.

Figure S5

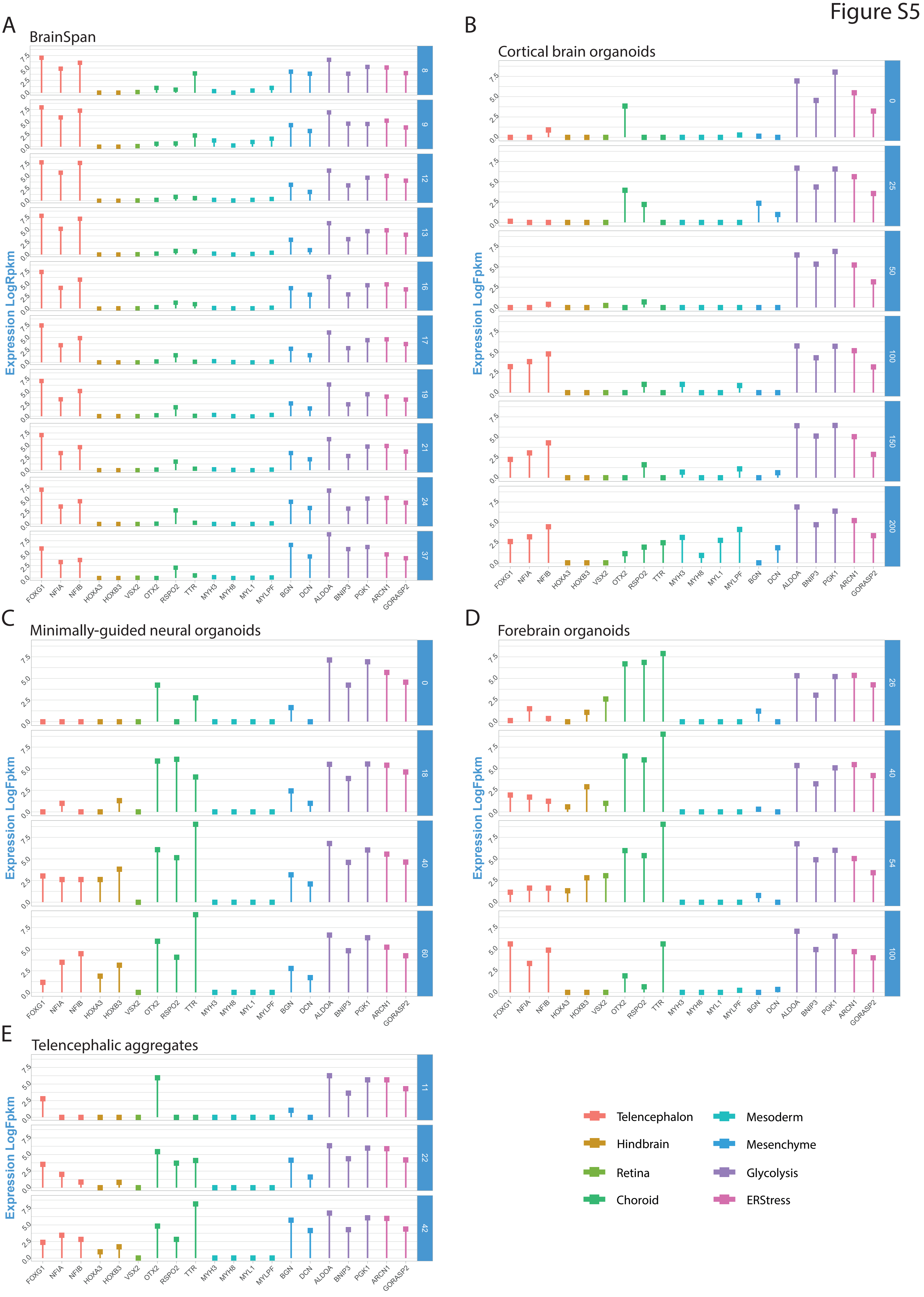
A
BrainSpan
Expression LogRpkm
8
9
12
13
16
17
19
21
24
37
B
Cortical brain organoids
Expression LogFpkm
0
25
50
100
150
200
C
Minimally-guided neural organoids
Expression LogFpkm
0
18
40
60
D
Forebrain organoids
Expression LogFpkm
26
40
54
100
E
Telencephalic aggregates
Expression LogFpkm
11
22
42
0.0
2.5
5.0
7.5
FOXG1
NFIA
NFIB
HOXA3
HOXB3
VSX2
OTX2
RSPO2
TTR
MYH3
MYH8
MYL1
MYLPF
BGN
DCN
ALDOA
BNIP3
PGK1
ARCN1
GORASP2
Telencephalon
Hindbrain
Retina
Choroid
Mesoderm
Mesenchyme
Glycolysis
ERStress

#### Supplementary Figure 5

Lollipops showing the expression levels of gene signatures related to brain areas, off-target tissues and cell stress in BS **(A)**, CBO **(B)**, MGO **(C)**, FO **(D)** and TA **(E)**. Expression levels (Log2Fpkm or Log2Rpm) along time-points are reported in each dataset as the mean value across replicates. Each bar colour corresponds to a specific signature, as reported in the plot legend.

Figure S6

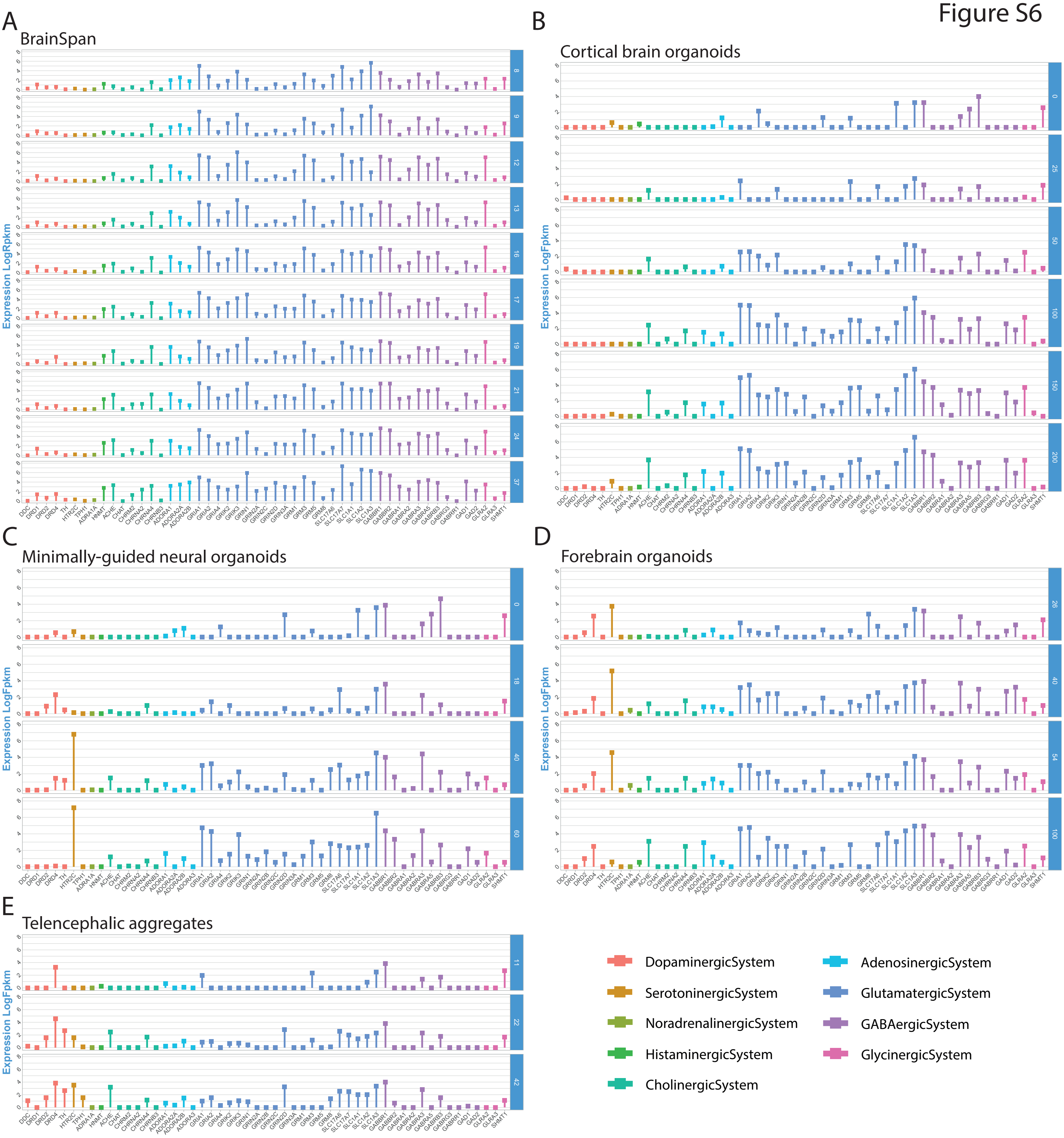
A
BrainSpan
Expression LogRpkm
B
Cortical brain organoids
Expression LogFpkm
C
Minimally-guided neural organoids
Expression LogFpkm
D
Forebrain organoids
Expression LogFpkm
E
Telencephalic aggregates
Expression LogFpkm
DopaminergicSystem
SerotoninergicSystem
NoradrenalinergicSystem
HistaminergicSystem
CholinergicSystem
AdenosinergicSystem
GlutamatergicSystem
GABAergicSystem
GlycinergicSystem

#### Supplementary Figure 6

Lollipops showing the expression levels of gene signatures related to neurotransmission systems in BS **(A)**, CBO **(B)**, MGO **(C)**, FO **(D)** and TA **(E)**. Expression levels (Log2Fpkm or Log2Rpm) along time-points are reported in each dataset as the mean value across replicates. Each bar colour corresponds to a specific signature, as reported in the plot legend.

Figure S7

A

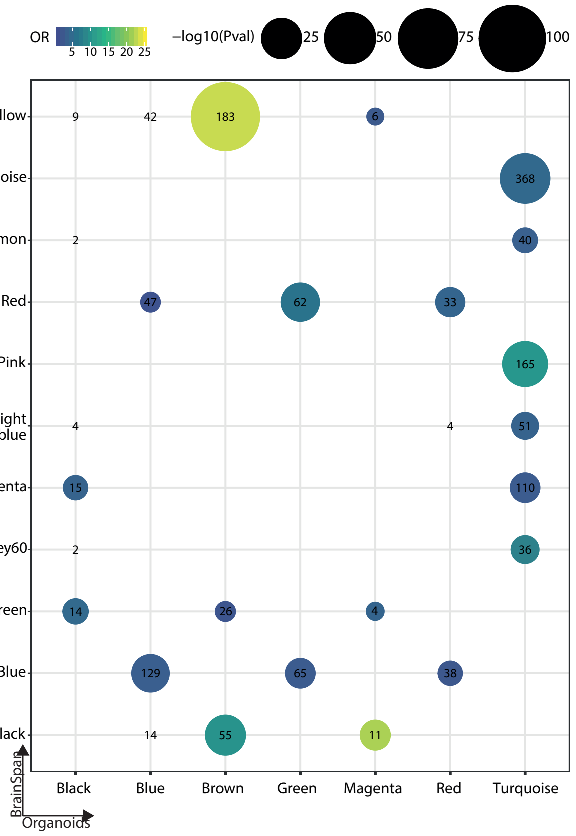

OR
5 10 15 20 25
−log10(Pval)
25
50
75
100
Yellow
Turquoise
Salmon
Red
Pink
Midnight blue
Magenta
Grey60
Green
Blue
Black
9
42
183
6
368
2
40
47
62
33
165
4
4
51
15
110
2
36
14
26
4
129
65
38
14
55
11
Black
Blue
Brown
Green
Magenta
Red
Turquoise
BrainSpan
Organoids

#### Supplementary Figure 7

Overlap between BrainSpan and Cortical Brain Organoid gene modules identified by WGCNA. Dot plot displaying the overlap between BS (y-axis) and CBO (x-axis) gene modules identified by WGCNA. Numbers represent shared genes and are shown for overlaps with odds ratio (OR) > 1, while dots are reported for those having also p-value $< 0.05$. Dot colour is assigned according to OR values and dot size according to p-value.

Figure S8

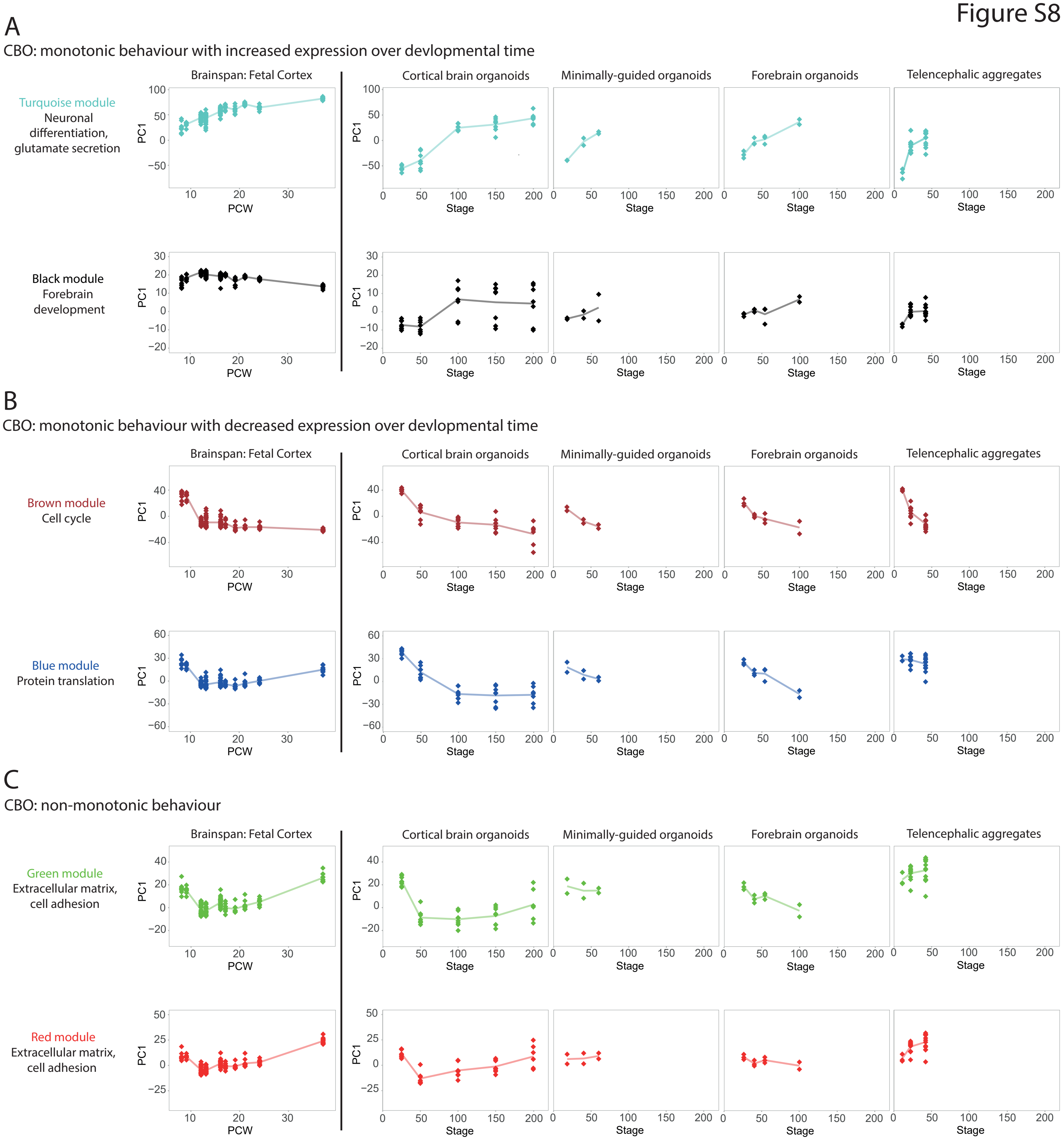
A
CBO: monotonic behaviour with increased expression over devlopmental time
Brainspan: Fetal Cortex
Cortical brain organoids
Minimally-guided organoids
Forebrain organoids
Telencephalic aggregates
Turquoise module
Neuronal differentiation, glutamate secretion
Black module
Forebrain development
PC1
PCW
Stage
B
CBO: monotonic behaviour with decreased expression over devlopmental time
Brown module
Cell cycle
Blue module
Protein translation
C
CBO: non-monotonic behaviour
Green module
Extracellular matrix, cell adhesion
Red module
Extracellular matrix, cell adhesion

#### Supplementary Figure 8

Behaviour of the module eigengene for Cortical Brain Organoids gene co-expression modules in fetal cortex and brain organoids datasets. **(A)** Visualization of CBO_Turquoise and CBO_Black eigengene module in prenatal fetal cortex as well as in the brain organoids' datasets. PCA was performed to calculate the module eigengene on the CBO dataset and then applied to BS prenatal cortex, as well as each of the other brain organoid datasets. Each dot represents a data point, while the line connects the median value for each post-conceptional week (BS) or differentiation day (BO). The same analysis and visualization are applied to CBO_Brown and CBO_Blue (**B**, decreasing during differentiation) and CBO_Green and CBO_Red (**C**, non-monotonic behaviour).
